# An infusible decellularized extracellular matrix material binds to vasculature in infarcted myocardium and induces pro-reparative gene expression following acute myocardial infarction through inherent avidity and bioactive signaling

**DOI:** 10.1101/2025.11.12.687915

**Authors:** Michael B. Nguyen, Alexander Chen, Van K. Ninh, Maxwell C. McCabe, Quincy Lyons, Colin Luo, Benjamin D. Bridgelal, Connor Uhre, Julian Cheng, Kate E. Reimold, Selena Cao, Kirk C. Hansen, Kevin R. King, Karen L. Christman

## Abstract

To treat acute myocardial infarction immediately after reperfusion, we previously engineered an intravascularly infusible decellularized extracellular matrix (iECM) biomaterial that exerts immunomodulatory and pro-reparative effects. However, the impact of the heterogeneous contents of iECM on infarct localization and downstream biological function is unknown. Using liquid chromatography, iECM is separated into a high molecular weight (MW) and low MW component. Mass spectrometry confirms compositional similarity, while biochemical assays and transmission electron microscopy highlight differences in biochemical features and structure, revealing a nanofibrillar high MW component and a globular peptide low MW. Quartz crystal microbalance studies show binding of each component to basal lamina ECM proteins and endothelial cell surface receptors under flow, demonstrating the specificity of ECM biomaterials to permeable vasculature. *In vivo*, the low MW component reduces vascular permeability, while neither component alone achieves the retention levels of complete iECM. Using single-nucleus RNA sequencing to probe bioactivity, both components elicited comparable angiogenic, immunomodulatory, and pro-reparative transcriptional programs. These findings illustrate that highly coupled materials and biological characterization uncover fundamental behaviors and properties of iECM biomaterials. Additionally, we show the unique binding behavior of iECM to the gaps of permeable vasculature, which could be exploited for future nanomaterial design.

## 1. Introduction

Ischemic heart diseases continue to be a significant health care burden worldwide, with an estimated 9 million deaths attributed to ischemic heart disease^[1]^. A major contributor to ischemic heart disease is acute myocardial infarction (AMI), which is characterized by the restriction of blood flow to the myocardium, often caused by the buildup of atherosclerotic plaque in the coronary arteries. This ischemic damage is characterized by cardiomyocyte death, endothelial dysfunction, increased vascular permeability, immune cell infiltration, and oxidative stress^[2]^. As a result of injury, cardiac function is impaired and the resulting compensatory adaptations in the weeks and months following AMI is known as negative left ventricular (LV) remodeling, which ultimately leads to heart failure^[3]^. The most common treatment for AMI is percutaneous coronary intervention, which opens the artery via balloon angioplasty^[1]^. This reperfusion is necessary to save the at-risk tissue; however, the reintroduction of oxygen rich blood further exacerbates the inflammation and is known as ischemia-reperfusion injury^[4]^.

For restoring cardiac function after MI, biomaterial treatments such as cardiac patches and hydrogels have been utilized to exert immunomodulatory effects and support tissue repair^[5]^. While cardiac patches have shown therapeutic effects in preclinical models^[6]^, they require invasive surgery to access the epicardium for implantation, bringing with it significant risk of complications. Similarly, the injection of biomaterials into the myocardium can mitigate inflammation and support cardiac repair^[7]^ but will be limited to sub-acute and chronic delivery timepoints clinically, as injection into the weakened myocardium at acute timepoints carries the risk of arrythmias and ventricular rupture^[8]^. As such, to treat MI and ischemia-reperfusion injury acutely, infusible biomaterials that are intravascularly delivered have seen increased popularity^[9]^. However, these materials require a method of localization to the injured myocardium to minimize off-target effects and improve treatment efficacy.

For many applications, decellularized extracellular matrix (ECM) biomaterials have seen great interest due to their inherent immunomodulatory and pro-reparative motifs derived from their source tissue^[10]^ as well as their cost effective production. To leverage these properties for reducing AMI induced negative LV remodeling in a minimally invasive manner, we recently developed an intravascularly infusible ECM material (iECM)^[11]^. Through further processing of pepsin digested ECM (Fig. 1a), iECM can be delivered immediately upon reperfusion without thrombus formation despite containing collagen-like fibril components^[11]^. iECM localizes to vasculature in areas of inflammation, reduces vascular permeability, exerts immunomodulatory effects, and improves cardiac function in both small and large animal models of AMI^[11]^. It also reduced inflammation and improved animal survival in a model of severe systemic inflammation^[12]^.

**Figure 1.**
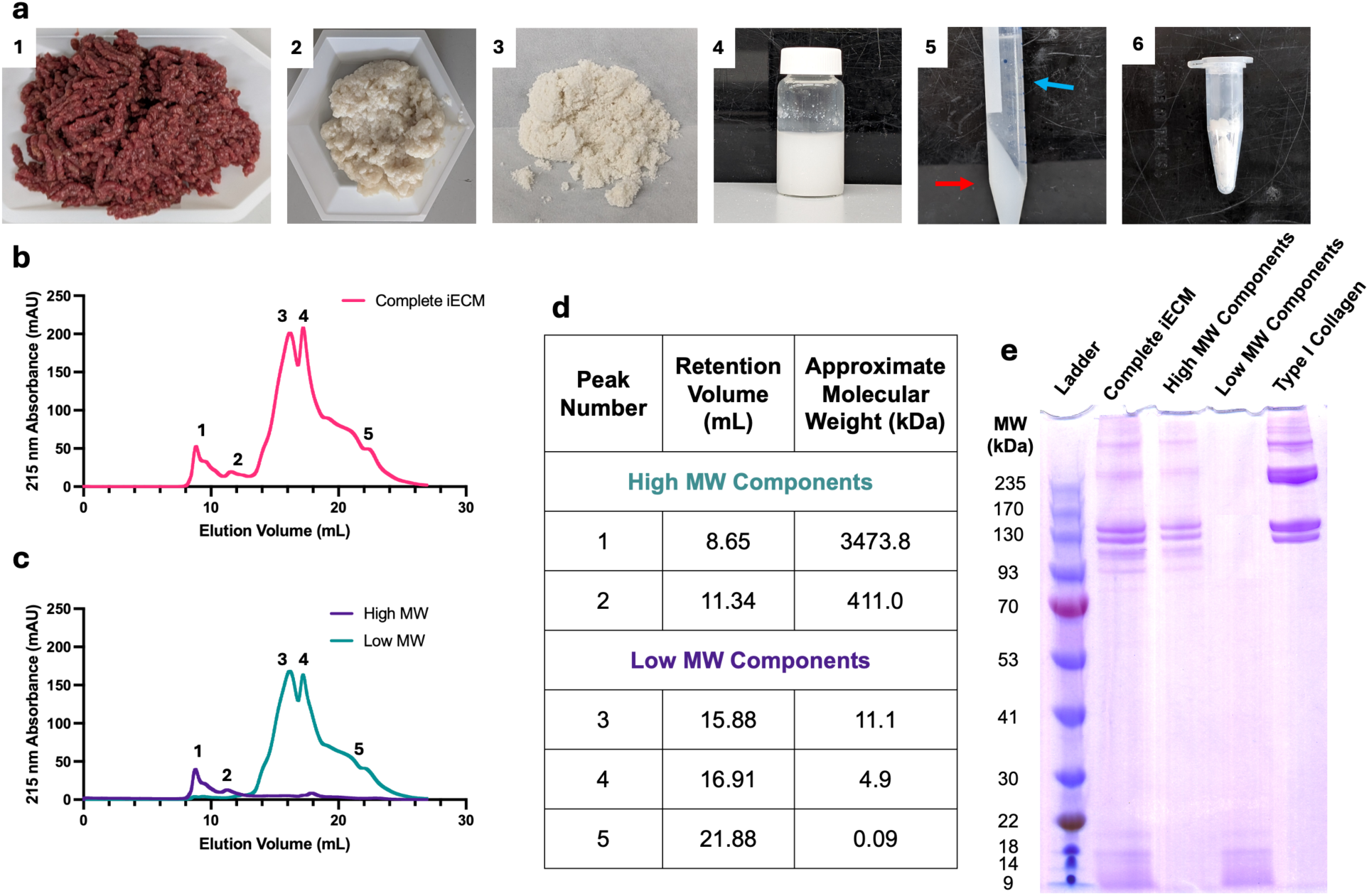
Characterization of iECM molecular weight distribution and separation of iECM components into two fractions. a) Fabrication of iECM by 1) mincing porcine LV tissue, 2) decellularizing with SDS, 3) lyophilizing and milling into a powder, 4) partially digesting with pepsin, 5) centrifuging to pellet out large particulate matter (red arrow), 6) sterile filtering the supernatant (blue arrow) and lyophilizing for storage. SEC traces of b) complete iECM and c) iECM separated into High MW components (blue) and Low MW components (purple). d) Quantification from SEC traces showing approximate molecular weight of iECM components. d) SDS-PAGE of complete iECM, High MW components, Low MW components, and collagen type I demonstrating efficient separation of iECM into two components.

While many studies have been done to determine how ECM materials work in the context of solid and hydrogel scaffolds, the mechanisms through which intravascularly delivered ECM materials interact with vasculature in inflamed tissue and exert therapeutic effects have yet to be understood. For synthetic nanomaterials, it has often be hypothesized that targeting of inflamed vasculature can be achieved through the use of anisotropic materials^[13]^ that more effectively marginate within the vasculature^[14]^. However, these materials often use a targeting moiety to bind to only a single component such as upregulated endothelial cell receptors that occur with inflammation like ICAM^[15]^ or the exposed underlying type IV collagen in the basement membrane in injured blood vessels^[16]^. Despite these rational targets, there has been no *in vivo* histological evidence showing these materials binding to the gaps in permeable endothelium. Given iECM was shown to have this unique localization^[11]^, we sought to investigate how iECM targets, which could inform the design of new nanomaterial therapeutics, as well as investigate the fundamental biochemical and biological behaviors of the material as part of our development of an efficacious minimally invasive treatment for AMI.

In this study, we applied size-based fractionation to separate iECM into its main components, enabling us to study how each component behaves in the context of inflammation and AMI. This method yielded two major components based on apparent molecular weight. We first characterized the morphological and biochemical properties to understand how each component would behave once delivered intravascularly. We then investigated the role of these components in reducing vascular permeability and inducing repair and regeneration of cardiac tissue via next generation sequencing. Altogether, this manuscript provides evidence for a new avenue for targeting and treating diseases associated with acute inflammation, leveraging both heterogeneous shape and specific high avidity binding as a mechanism for targeting and treating permeable vasculature.

## 2.Results and discussion

### 2.1: iECM consist of a high molecular weight nanofibrillar component and a soluble, low molecular weight peptide component

iECM was fabricated using decellularized porcine LV myocardial tissue similar to previous reports^[11]^ (Fig. 1a). Degree of decellularization was measured using a double stranded DNA (dsDNA) assay showing less than 5 ng dsDNA per mg of ECM as well as sufficient sodium dodecyl sulfate (SDS) removal (Fig S1a-b)^[17]^. Via SDS-PAGE, we saw that iECM was qualitatively consistent with previous batches (Fig. 1e)^[11, 18]^. Overall, this suggested that the iECM fabrication process is robust and reproducible.

To quantify the relative composition of iECM, we analyzed iECM with size exclusion chromatography (SEC) (Fig. 1b). The processing of iECM with 0.22 μm filtration and a viscosity comparable to saline^[11]^ enabled SEC analysis of the entirety of iECM, in contrast with hydrogel formulations of decellularized ECM^[19]^. The mobile phase was phosphate buffered saline to simulate the salt conditions that iECM would experience after intravascular delivery. We found that iECM followed a bimodal distribution comprised of a high molecular weight (High MW) component (MW > 100 kDa) and a low molecular weight component (Low MW) (MW < 10 kDa) (Fig. 1b). Based on 215 nm absorbance^[20]^, high MW components made up approximately 15% of the total iECM composition, while the low MW components constituted the remaining 85%. To separate out the two components based on MW, we leveraged the differences in solubility. The low MW components could be dissolved using 50% ethanol in water while the high MW components precipitated out (Fig. 1c-d). After separation and weighing, the percentages of each component was found to be approximately 19% and 81% high MW and low MW by mass, with this difference due to differences in the amino acid composition^[20]^ and the presence of non-protein components in the high MW component that did not appear on SEC. Separation of iECM components was confirmed via SEC and SDS-PAGE, where the peaks and bands of iECM components matched well to complete iECM, confirming that the conformation of the components was not affected by the separation process (Fig. 1c, e). Overall, these data show that the complete iECM can be efficiently fractioned into its 2 predominant components and studied independently.

To investigate the composition, label free quantification mass spectrometry (Fig. 2a-b, SI Table 1-2) showed that the high MW component was predominantly fibrillar collagens while the low MW component had a greater percentage of nonfibrillar components. We then compared the overlap in number of identified parent proteins and peptide sequences without factoring in relative abundances. Overall, 94 ± 3.5% of the identified parent proteins in the high MW were found in the low MW and 91 ± 0.1% of the identified parent proteins in the low MW were found in the high MW. For peptide sequences, 71 ± 2.3% of peptide sequences identified in the high MW were found in the low MW and 76 ± 0.6% of peptide sequences identified in the low MW were found in the high MW. This showed that both components were mostly derived from the same parent proteins. The lower overlap percentage in the peptide sequences is likely due to differences in protein abundances where a protein with higher abundance will generate more sequence coverage. Overall, the high MW and low MW components are compositionally similar, and the relative abundance is the primary difference. To investigate morphology, negative stained transmission electron microscopy (nsTEM) revealed that the high MW component contained fragmented, flexible nanofibers (Fig. 2c), similar to what was previously seen with complete iECM^[11]^. The low MW component contained globular peptides (Fig. 2d). Together these data suggest that the low MW component contains the products of pepsin digestion while the high MW component contained the partially digested nanofibers.

**Figure 2.**
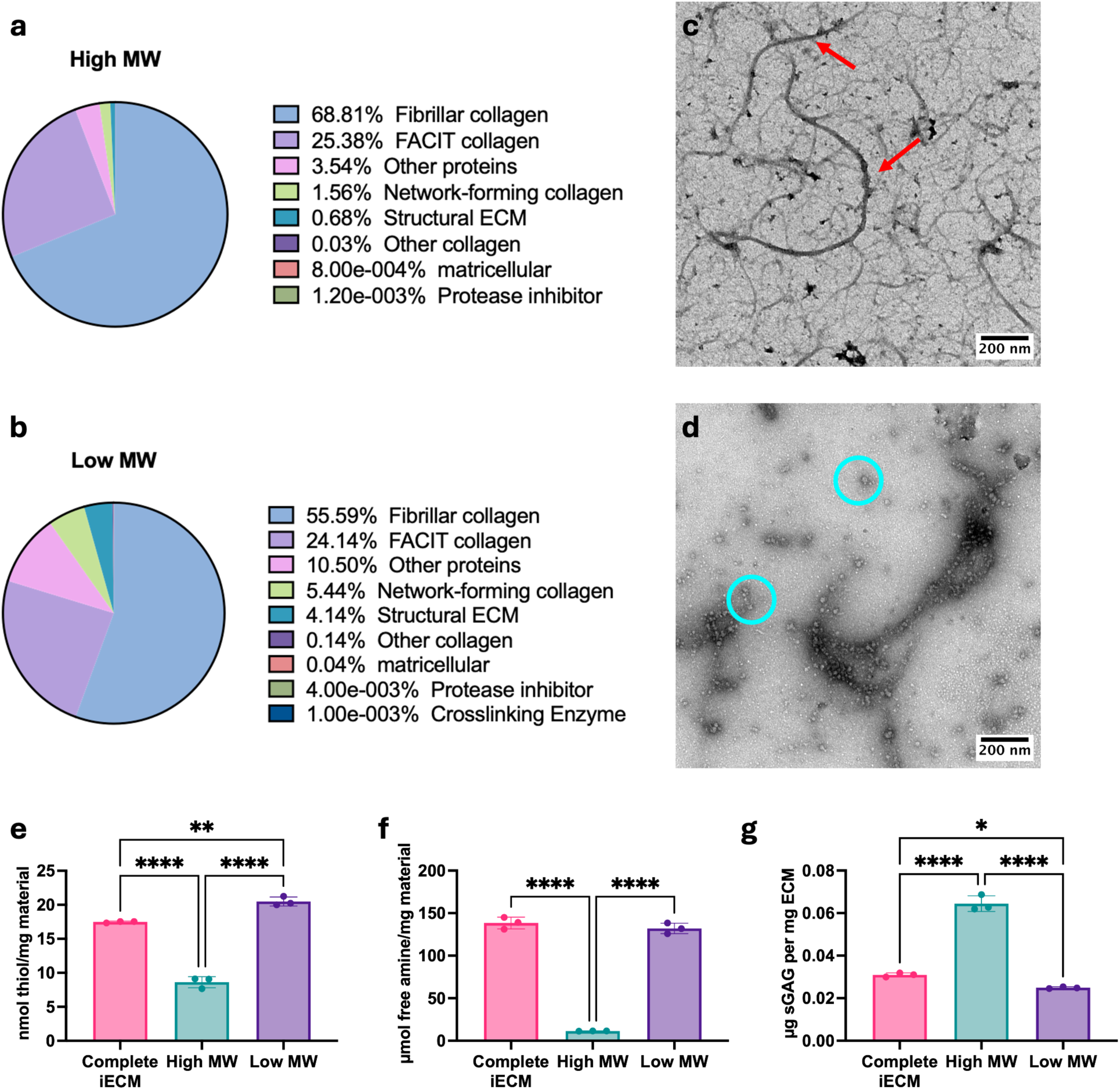
Compositional, morphological and biochemical analysis of iECM and its components. The fractions were analyzed via label free quantification of trypsin digested mass spectrometry for protein content with the relative spectral intensity shown for the a) High MW components and b) Low MW components. The morphologies were also visualized via TEM for morphology: c) high MW and d) low MW. Biochemical assays measuring the e) thiol content, f) free amine content and g) sGAG content of iECM, low MW and high MW components. Red arrows denote nanofibrils and cyan circles denote globular and nanofibril peptides. Only p < 0.05 displayed, * < 0.05, ** < 0.01, *** < 0.001, **** < 0.0001.

We next characterized the biochemical differences between the components. The free thiol component was significantly greater within the low MW component. Previous findings determined that iECM was capable of acting as an antioxidant^[7c, 11]^, with these data suggesting that potential reactive oxygen species sequestration is likely driven by the low MW component (Fig. 2e). We also found that the low MW component contained significantly more free amines (Fig. 2f) compared to the high MW component. This was expected given the hypothesis that the low MW was comprise predominantly of digested ECM peptides, as the cleaving of ECM proteins into peptides would generate more free amines through the breakage of amide bonds. Finally, we found that the high MW component contained most of the sulfated glycosaminoglycans (sGAG) (Fig. 2g). sGAGs, typically found as part of proteoglycans, are desirable within decellularized ECM materials for their bioactivity^[17]^. Therefore, based on the biochemical readouts, both components may contribute a unique mechanism to the overall biological activity of the complete iECM.

### 2.2 iECM and its components specifically bind to extracellular matrix proteins and cell surface receptors

We next investigated how these observed differences translated to the binding capabilities of iECM components. Since iECM was found to colocalize with permeable vasculature post-MI^[11]^, we evaluated binding to underlying ECM components exposed with permeable vasculature such as laminins^[21]^, type IV collagen^[21]^, and type I collagen^[22]^. Additionally, endothelial cell damage and dysfunction caused by MI results in upregulated integrin α1β1^[23]^, intracellular adhesion molecule 1 (ICAM/CD54)^[24]^, and P-selectin^[25]^. Quartz crystal microbalance (QCM) is sensitive to surface binding under flow and has previously been used to evaluate peptide binding to ECM surfaces^[26]^ and antibodies to laminin surfaces^[27]^. Since the moles of the iECM components cannot be determined due to the polydispersity, we evaluated relative binding efficiency for a target by delivering the high and low MW at their respective proportions in complete iECM. Using previous protocols^[28]^, ECM proteins and cell receptor proteins were conjugated to functionalized lipids. Since the lipid-protein coating had a high dissipation, it was viscoelastic and thick. Thus, the lower overtones more accurately represent the binding to the conjugated proteins at the top of the thick layer^[26]^. We reported the third overtone frequency shift as previously done for hydrated laminin ^[27]^ and other QCM results^[29]^. Using antibodies to validate target coating, as was investigated for laminin^[27]^, we found varying degrees of antibody binding (Fig. S2a) due to the variation in the efficiency of ECM and cell receptor protein conjugation. Nonetheless, our antibody binding to laminin was similar to previous results^[27]^, giving confidence in successful conjugation.

Overall, the high MW component had increased binding to all of the coatings except for type IV collagen and P-selectin (Fig. 3a-b). This suggested that the high MW had the greatest specificity to the various components exposed in the permeable vasculature. Interestingly, the high MW binding for laminin 111, type I collagen, ICAM, and integrin α1β1 was significantly higher than both complete iECM and low MW, which suggested interactions between the components within complete iECM and therefore allowing the high MW to bind more efficiently when on its own. To assess the interactions of the components, we coated the surface with the high MW or low MW component and flowed the opposite component. Overall, we found that the high MW bound more to the low MW (Fig S2b). This difference in binding was because the low MW had significantly more free amines (Fig 2f) and therefore increased conjugation efficiency to the lipid surface allowing for a greater degree of binding sites. Overall, these data suggest, the two components interact with each other, which may impact the iECM localization *in vivo*.

**Figure 3.**
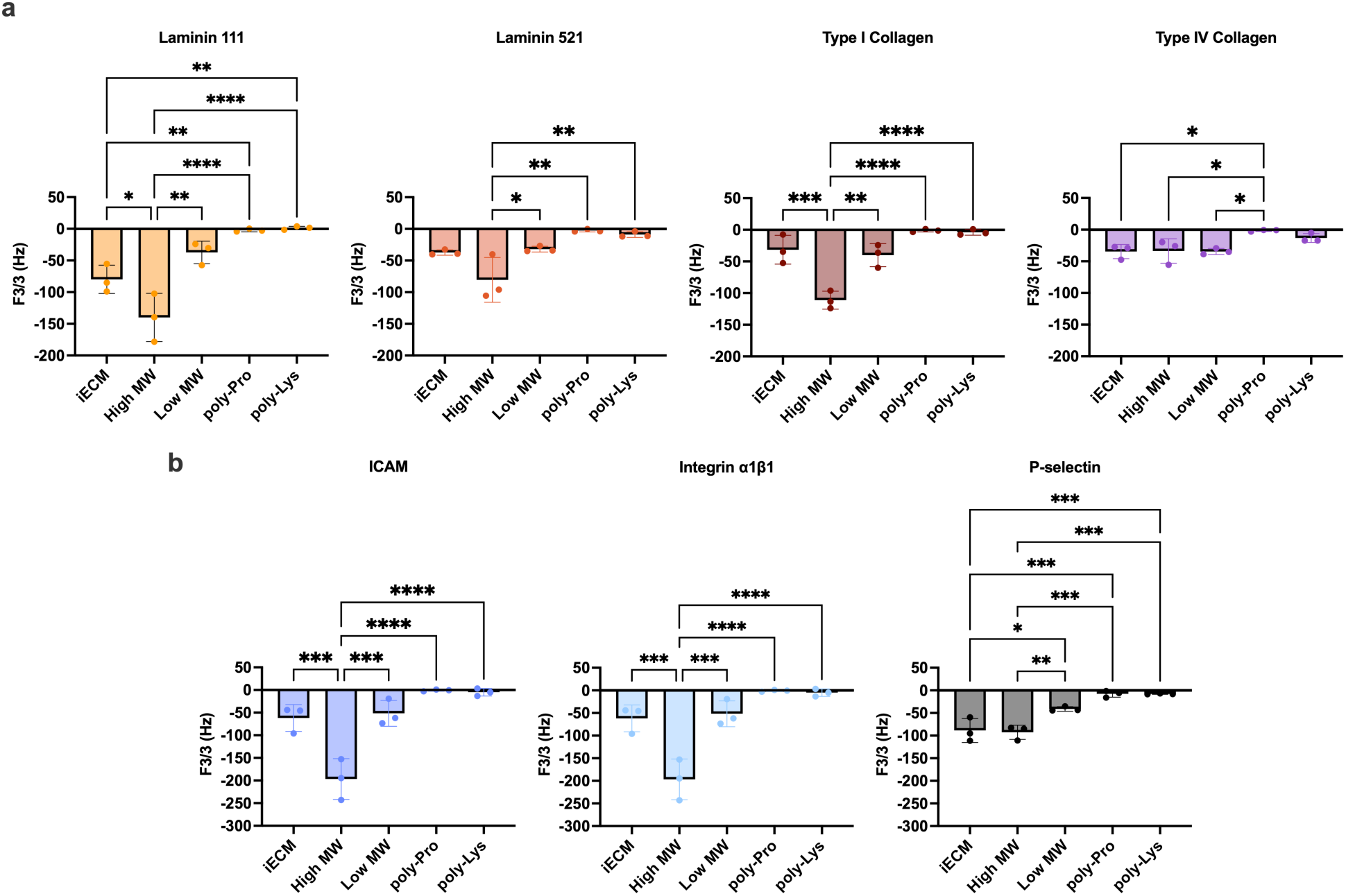
Evaluation of binding of iECM and its components to immobilized ECM proteins and cell receptors via quartz crystal microbalance. Display of the average 3^rd^ harmonic frequency shift values with standard deviations to show variance for binding to a) ECM components and b) cell surface receptors. Lower frequency represents greater binding of the flowed iECM or components to the immobilized protein of interest. Comparisons were only made between the same chemical conjugated coating. One-way ANOVA with Tukey’s post-hoc between plotted groups. *p < 0.05, **p < 0.01, ***p < 0.001, ****p < 0.0001.

As controls, poly-L-proline (neutral) and poly-L-lysine (positive) with similar MWs to the low MW component were used. The poly-L-lysine had a larger shift in frequency compared to the poly-L-proline suggesting an improvement of binding with an electrostatic charge. However, relative to the iECM and its components, the frequency shifts were significantly lower for most targets (Fig. 3). This suggested that complete iECM and its components have specific binding to the various ECM and cellular receptors, therefore confirming the previously observed colocalization with the gaps of permeable vasculature as specific binding. This is not surprising given that self- and cross-binding between ECM proteins is well known, both in the context of native tissue and solid biomaterial development^[30]^. Our data demonstrates that the distinct components of iECM retain the unique binding behavior to both ECM proteins and cell surface receptors while under flow, enabling them to target multiple aspects of endothelial dysfunction. The findings highlight a unique behavior of iECM with varying binding affinities to other ECM components and cellular receptors and illustrate the advantage of diverse specific binding modes for a heterogeneous ECM biomaterial.

### 2.3 : iECM components readily localize to sites of permeable vasculature and reduce vascular permeability in vivo

Following material characterization of the iECM components, we investigated their biodistribution and retention in a rat model of MI to determine which component may drive localization and retention in inflamed vasculature *in vivo*. Following MI, a simulated intracoronary injection was performed with fluorescently tagged complete iECM, high MW, and low MW components at the relative mass-based concentrations (10 mg/mL complete iECM, 1.9 mg/mL high MW components, 8.1 mg/mL low MW components). To match the relative fluorescence of each of the iECM components, iECM was fluorescently labeled prior to separation by ethanol. Hearts were excised and fluorescently imaged at 1-hour and 3 days post-MI. When images of all scanned tissues are adjusted to the same dynamic ranges, the localization of complete iECM and low MW were more visible compared saline and high MW (Fig. S3a). However, when the high MW and saline were adjusted alone, localization of the high MW was apparent (Figure S3b). This was because of differences in fluorophore conjugation efficiency due to significant differences in free amine content between the components. While the high MW component was approximately 19% by mass of iECM, it only contained 2.6% of the fluorescent dye (Fig. S3c). Similarly, the low MW was 81% by mass of iECM and 97.4% of the fluorescent dye (Fig. S3c). Nonetheless, when evaluating total signal, complete iECM signal was greater than both the high MW and low MW components combined at 1 hour (Fig 4a), suggesting improved retention when delivered together. At day 3, the complete iECM still had more signal than both components further supporting improved retention when the components are delivered together (Fig. 4b). After normalization to account for the differences in fluorophore conjugation efficiency, we found that the high MW components localized to the infarct significantly more than the complete iECM or the low MW components at 1 hour and 3 days (Figure 4c-d). These data are in alignment with the QCM data that suggested greater efficiency to our selected substrates by the high MW components alone compared to the low MW components. Additionally, seeing the significantly higher high MW normalized signal at 3 days suggested that the high MW component takes longer to degrade and clear from the infarct, potentially due to its macromolecular structure. These data suggest that the high MW exhibits the greatest binding efficiency and retention *in vivo,* informing future iECM formulations and bioinspired infusible biomaterial design.

**Figure 4.**
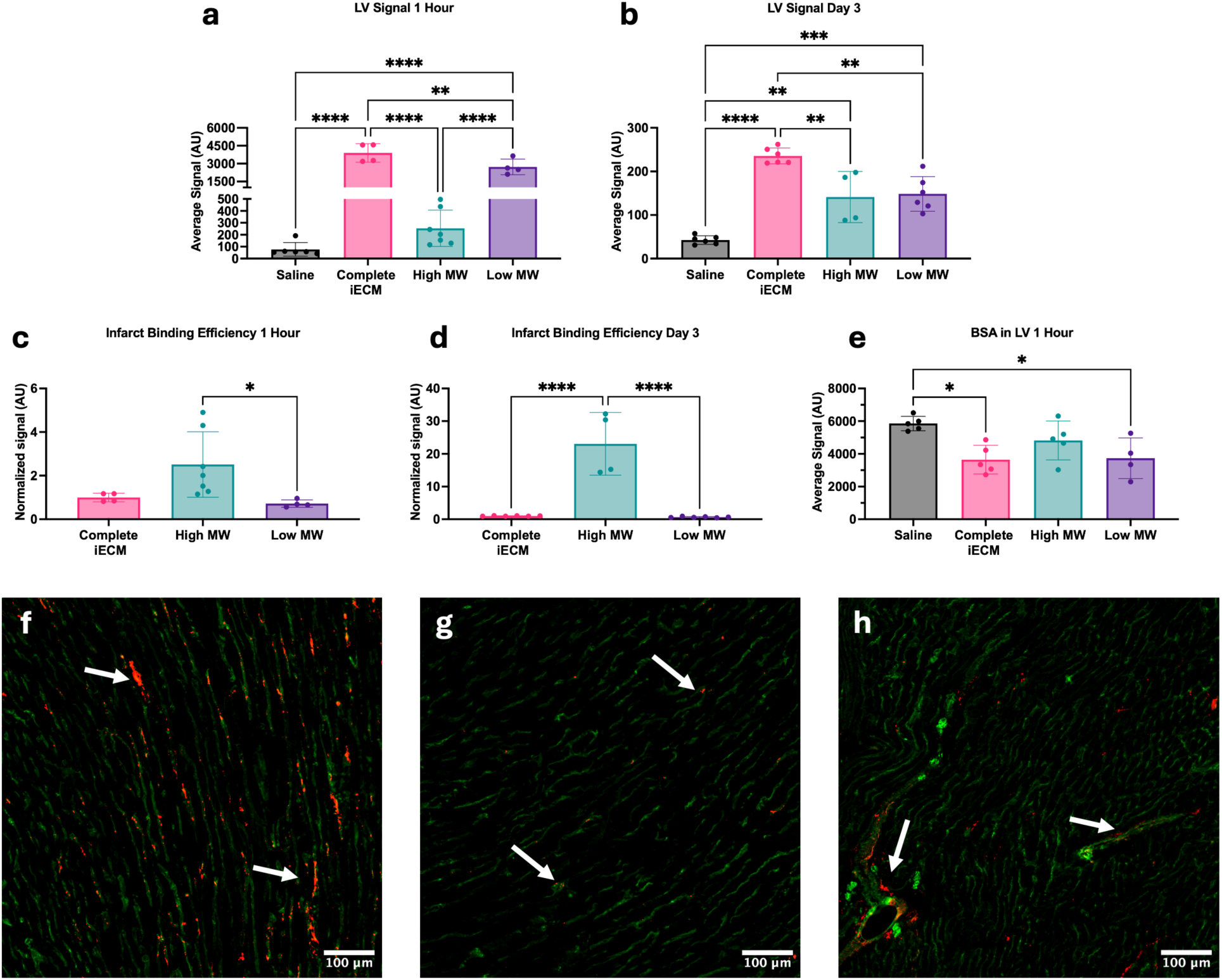
Biodistribution, retention and effect of the iECM and its fractions on vascular permeability 1 hour and 3 days post-MI. Quantified fluorescent signal of iECM and iECM fractions a) 1 hour post reperfusion and b) 3 days post reperfusion. The LV retention signal normalized to the fluorophore conjugation efficiency for complete iECM and its fractions at c) 1 hour and d) 3 days. e) Vascular permeability of infarct as determined by residual BSA in LV at 1 hour. Immunofluorescent stains of the f) iECM, g) high MW, and h) low MW in red and isolectin for vessels in green within the infarct. White arrows denote example points of colocalization with vasculature. One-way ANOVA with Tukey’s post-hoc. *p < 0.05, **p < 0.01, ***p < 0.001, ****p < 0.0001.

Post-MI, inflammation from prolonged ischemia and reperfusion induces endothelial dysfunction, resulting in increased vascular permeability, edema, and additional tissue damage^[31]^. Previous work demonstrated that iECM reduced vascular permeability following infusion, which is a hypothesized contributor to iECM’s therapeutic effects^[11]^. Using a fluorescently tagged bovine serum albumin (BSA) tracer injected 30 minutes post reperfusion, both complete iECM and the low MW components alone were able to significantly reduce vascular permeability compared to the saline control (Fig. 4e). Despite high binding efficiency, high MW components were not significantly different compared to any group (Fig. 4e). The was likely due to the dosing, as the relative mass of high MW delivered is approximately one fourth of the low MW. Via fluorescent staining, we saw colocalization of iECM, high MW, and low MW with endothelial cells within the infarct (Figure 4f-h), confirming their localization to the infarcted vasculature similar to previous results^[11]^. Overall, these data showed that the iECM components bind to the permeable infarct vasculature. While the low MW components alone were able to reduce vascular permeability soon after reperfusion, the high MW had significantly higher binding efficiency within the infarct at both 1 hour and 3 days post-MI. Altogether, the complete iECM exhibited the highest total signal at both 1 hour and 3 day, suggesting an improved binding and retention over the individual components.

### 2.4 : Delivery of iECM and its components result in transcriptomic changes following AMI

Previous work with iECM has demonstrated its ability to induce transcriptomic changes within the infarct^[32]^. Given that both components appeared to localize to the infarct, we then sought to better understand the role of each of iECM’s components on its bioactivity. To this end, we performed single-nucleus RNA sequencing (snRNAseq) on tissue samples isolated from animals treated with complete iECM, iECM components, or saline 7 days post-MI since previous work has shown that iECM exhibits the most abundant differences in gene expression at this timepoint^[32]^. Infarcts from each treatment were quantified to ensure sufficient injury to the myocardium before nuclei isolation; infarct sizes were not significantly different between treatments (Figure S4a-c). iECM and saline datasets from the previous snRNAseq investigation^[32]^ were included to increase statistical robustness. The SoupX function was used to remove ambient RNA contamination, and the features and counts were evaluated (Fig. S5a-b). The datasets from all treatments were integrated and coarsely clustered. We identified fibroblasts, cardiomyocytes, endothelial cells, macrophages, mural cells (pericytes and vascular smooth muscle cells), proliferating cells, T cells, B cells, endocardial cells, neuronal cells, lymphatic endothelial cells and mesothelial cells (Fig S5c-d) using marker genes previously found in literature^[32]^. The relative proportions of each cell type are also shown (Fig S5c) and outlined by replicate in SI Table 3. After separating the data by cell type, the data was also split by treatment. The saline data was the normalized reference for integration for each treatment. Integration was separate between treatments so that the cells from the other treatments would not impact the integration and bioinformatic removal of contamination and doublets. After contamination and doublet removal, the entire cluster was evaluated for differentially expressed genes (DEGs) between the saline and treatment groups (SI Table 4-12). The upregulated and downregulated DEGs were then analyzed via gene ontology (GO) pathway enrichment for biological processes, molecular functions and cellular components.

#### 2.4.1 iECM and its components are robustly angiogenic

Within the endothelial cell (EC) population (*Flt1^+^/Cdh5^+^)* (**Fig. 5a-c, S6a-c**), iECM upregulation of genes such as *Notch1* and *Plxnd1* (Fig 5a) suggested an angiogenic response, which matches previous findings^[32]^. Previous reports have shown that *Notch1* signaling is closely tied to the expression of cardiac markers in circulating progenitor cells by aiding in the epithelial-to-mesenchymal transition^[33]^. When looking at the high MW, significant signs of angiogenesis and endothelial progenitor cell development can be seen (Fig 5b). Upregulation of *Tgfbr1*, *Flt1, Plxnd1, Nrp1,* and *Nrp2* further support angiogenesis*^[34]^*. There is integrin upregulation via *Itga1* and *Itga6*, which aids adhesion and migration of circulating cells, chemoattraction via upregulated *Cxcl12*, and differentiation into endothelial progenitors evidenced with upregulated *Cd34* and *Kdr (Vegfr2)*. The low MW also shows similar signs of angiogenesis via *Flt1*, *Itga6*, *Pecam1*, and *Nrp1* (Fig 5c). Overall, these data suggest that the high MW shows evidence for endothelial progenitor recruitment while the low MW only exhibits a robust angiogenic response.

**Figure 5.**
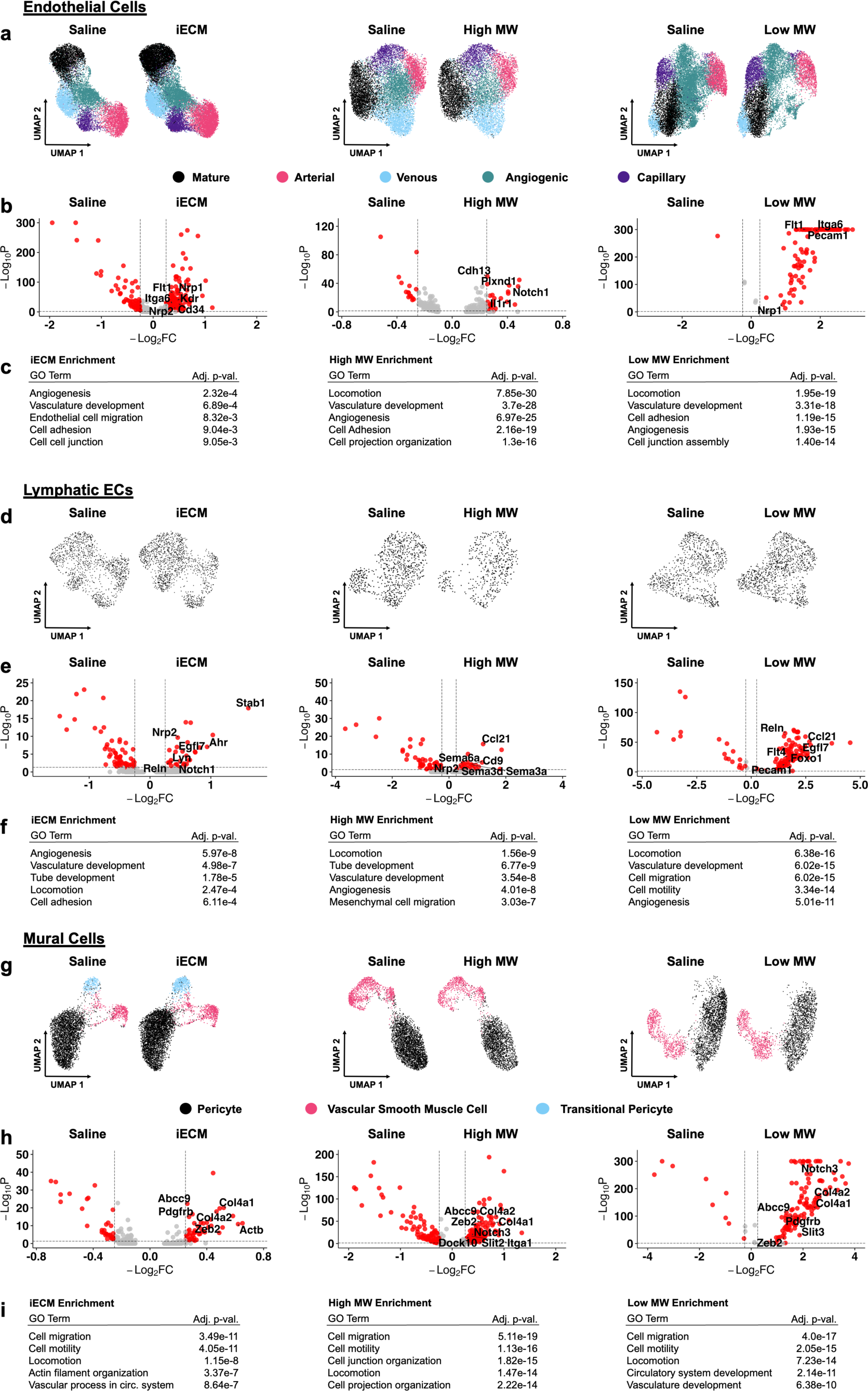
Single nucleus RNA sequencing integration, differential gene expression and gene ontology enrichment analysis of endothelial cells, lymphatic endothelial cells and mural cells for iECM, high MW and low MW compared to saline treatment. The UMAPs, volcano plot and enriched GO terms for iECM, high MW and low MW vs. saline for a-c) endothelial cells, d-f) lymphatic endothelial cells and g-i) mural cells suggest that all treatments resulted in angiogenic and vasculature development related responses.

Within the lymphatic EC population (*Prox1^+^/Flt4^+^I*) (Fig. 5d-f), iECM treatment showed upregulation of *Notch1* and *Egfl7,* which are angiogenic markers^[35]^ matching previous work^[32]^. Moreover, iECM treated lymphatic ECs also exhibit *Reln*, a lymphoangiocrine that is tied to cardiac growth and repair^[36]^, and *Ahr* which has shown to improve cardiac function^[37]^. Within the high MW group, there was via SEMA signaling and upregulation of *Ccl21*, *Nrp2*, and *Cd9* (Fig 5d), which showed a similar angiogenic response in addition to an immunomodulatory response and mesenchymal cell migration^[38]^. Within the high MW group, there was via SEMA signaling and upregulation of *Ccl21*, *Nrp2*, and Cd9 upregulation (Fig 5e), which showed a similar angiogenic response in addition to an immunomodulatory response and mesenchymal cell migration^[38]^. Within the low MW (Fig. 5f), angiogenesis was also seen via *Egfl7* in addition to the lymphoangiocrine *Reln*. There was also robust SEMA signaling and immune cell recruitment via *Ccl21*, *Flt4*, and *Pecam1* and upregulation of *Foxo1* suggesting mesenchymal cell recruitment^[39]^. The lymphatic EC response with iECM treatment matched the angiogenic response of the endothelial cells. The high and low components both showed robust evidence of angiogenesis in addition to signs of immunomodulation, lymphoangiocrine signaling, and mesenchymal cell recruitment.

Within the mural cell subset, which includes pericytes and vascular smooth muscle cells (*Abcc9^+^* or *Myh11^+^*) (Fig. 5g-i, S7a-c), all treatments generated upregulation of *Col4a1* and *Col4a2* (Fig 5g-i), which are components of the basement membrane of vasculature^[40]^.^[40]^. This suggested that mural cells were supporting the stabilization of the vasculature within the infarct due to treatment with iECM and its components. In addition, all treatments showed upregulation of *Zeb2*, which has been linked with cardiomyocyte growth and promotion of angiogenesis^[41]^, and upregulation of *Abcc9* which has been linked with smooth muscle cell development^[42]^. The high and low MW both showed upregulation of *Notch3* further supporting smooth muscle cell development^[43]^ and *Dock10*, which has been linked with regulation of cardiac function under stress^[44]^. The low MW also had *Slit3* upregulation, which has been tied to mesenchymal cell migration and vasculature development^[45]^. Conversely, all treatments also showed upregulation of *Il34*, which has been linked with macrophage recruitment and increased inflammation post-MI^[46]^. The high and low MW also showed *Rock1* upregulation, which revealed signs of reduced endothelial cell development^[47]^. The high MW also showed *Slit2* upregulation which has also been linked with reduced endothelial cell development^[48]^. Overall, these findings suggest that the mural cells from all treatments show clear signs of vasculature support and maturation. The angiogenesis inhibiting gene expression is likely tied to maturation of the newly formed vessels seen in the endothelial response. Moreover, *Slit2* has been found to suppress endothelial cell development yet also reduce the inflammatory response post-MI.

When the vascular cell populations are considered as a whole, iECM and its components exhibit strong signs of angiogenesis, vessel development, and maturation. Particularly, the high MW exhibits the most robust signs of endothelial progenitor cell recruitment. In addition, evidence points to immunomodulation towards cardiac repair and the recruitment and signaling for migrating mesenchymal cells. Within each treatment, a variety of subpopulations were identified. These populations were generally conserved across treatment (Fig. S6-9), which suggests that the therapeutic effects of the individual iECM components was not significantly different.

#### 2.4.2 iECM and its components show signs of immunomodulation

The major immune cell types captured included B cells (*Cd74^+^/Bank1^+^)*, T cells (*Themis^+^/Bcl11b^+^)* and macrophages (*Mrc1^+^/Csf1r*^+^ or *Maf^+^/Abca1^+^)* (Fig 6, S8-10). While neutrophils have been widely implicated in ischemia reperfusion injury^[2b]^ and have been a target of interest for immunomodulation post-MI, they were not studied here since single nucleus RNAseq does not capture neutrophils, but is required for cardiomyocytes. Furthermore, neutrophils peak at 24 hours post-MI and are reduced by day 7^[2b]^.

**Figure 6.**
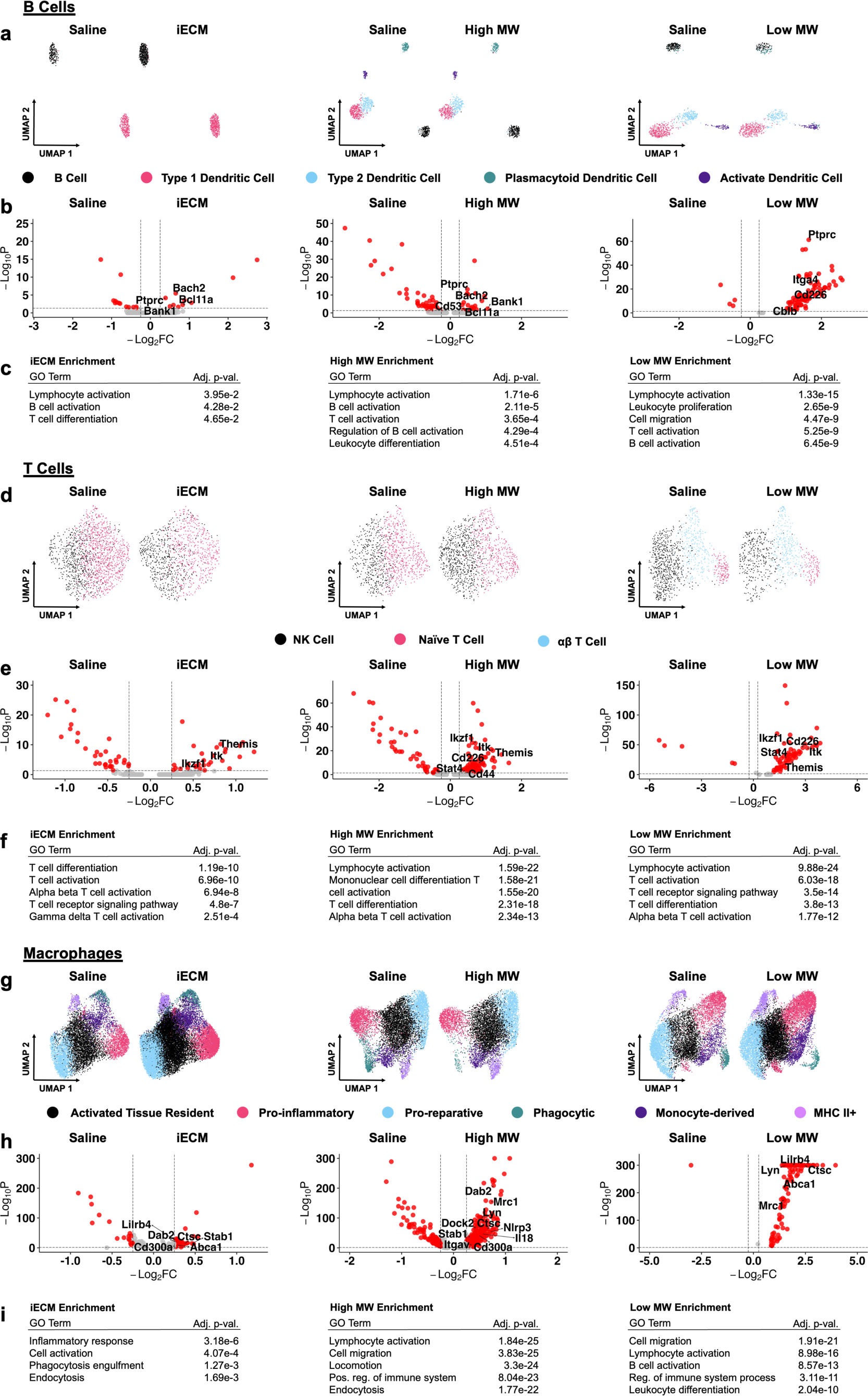
Single nucleus RNA sequencing integration, differential gene expression and gene ontology enrichment analysis of B cells, T cells, and macrophages for iECM, high MW and low MW treatment compared to saline treatment. The UMAPs, volcano plots and enriched GO terms for iECM, high MW and low MW vs. saline for B cells (a, b, c), T cells (d, e, f) and macrophages (g, h, i) respectively. All treatments generated similar responses versus saline via activation of the immune cell type of captured. Overall, the M2-macrophage response was sustained at day 7 by all treatments vs. saline.

The B cell population contained other antigen presenting cells like dendritic cells (Fig 6a-c, S8). There were significant B cell activation and development in all treatments via *Bcl11a* and *Ptprc* upregulation. In addition, there was *Bank1* expression for iECM and high MW groups and *Cd53* in the high MW (Fig S6b) further supporting activation^[49]^. The B cells of iECM and high MW also showed upregulation of *Bach2*, which has been linked to reduced cardiac hypertrophy^[50]^. The low MW also showed signs of B cell development via *Bcl11a*, *Ptprc*, *Itga4* and *Cd226*. There was also *Cblb* upregulation suggesting a potential reduction in inflammatory cascades^[51]^.

The T cell population, which also contained an NK cell subpopulation, in all treatments suggested similar activation and differentiation via upregulation of *Themis*, *Bcl11b*, and *Ikzf1* (**Fig 6d-f, S9**). Additionally, the high MW exhibited *Cd44* upregulation, which has been linked with infarct healing via immunomodulation^[52]^ as well as *Stat4*, which is linked to alleviation of injury post-MI^[53]^. Despite this, a pro inflammatory T cell phenotype existed seen through *Itk* expression^[54]^ across all treatments and *Cd226* expression^[55]^ in the high and low MW. Overall, this data suggested that while iECM and its components do not fully ablate the inflammatory response post-MI, all treatments generated signs of an immunomodulation toward a pro-repair response.

Within the macrophage subset (**Fig 6g-i, S10a-c**), there was a mixed expression of proinflammatory M1-like and pro-reparative M2-like macrophages. For iECM treatment (Fig. 6g), pro-reparative genes^[56]^ are expressed like *Stab1*^[56a]^ and *Dab2*, alongside pro-inflammatory genes^[57]^ like *Lilrb4*, *Abca1*, *Cd300a* and *Ctsc*. In the high MW treatment (Fig. 6h), upregulation of *Lilrb4, Nlrp3*, *Tlr7*, *Il18*, *Dock2*, *Ctsc*, *Cd300a*, and *Abca1* signified an M1 macrophage presence^[57]^ while upregulation of *Mrc1*, *Dab2*, *Stab1*, *Itgav, Lyn*, and *Csf1r* supported a significant M2-macrophage population^[32]^. The low MW treatment (Fig 6i) showed M1 gene expression via *Ctsc*, *Lilrb4* and *Abca1* as well as M2 gene expression via *Lyn* and *Mrc1* (Fig 6i). Previous findings showed that iECM accelerated and sustained the M2-macrophage response from day 1 through day 7 post-MI^[32]^. These data further support that claim and that both components also exhibit a similar sustained pro-reparative macrophage phenotype at day 7.

#### 2.4.3 iECM and its components promote a pro-regenerative phenotype

Within the cardiomyocyte subset (*Ryr2^+^/Tnni3k^+^*) (**Fig. 7a-c, S11a-c**), the overall phenotype across treatments suggests a shift towards cardioprotection, although some fibrotic and hypertrophic response still exists. With iECM treatment (Fig. 7a), upregulation of *Vegfa Ddah1*, *Srsf3, Ndrg4,* and *Ddx17*, suggest cardiac angiogenesis and preservation of systolic function^[58]^. Conversely, *Rnf207* was also upregulated with iECM treatment, which has been tied to cardiac hypertrophy^[59]^. With high MW treatment (Fig. 7b), there was upregulation of *Erbb4,* which has been linked with mesenchymal cell myocardial repair^[60]^. In addition, the high MW had upregulation of *Corin, Tbx20*, *Pde3a*, and *Nrp1*, which have been linked with reduced cardiomyocyte apoptosis^[61]^ and upregulation of *Fgf12* and *Fgf13*, which has been linked with reduced cardiac remodeling and reduced inflammation^[62]^. With low MW treatment (Fig. 7c), similar to the high MW, there was also upregulation of *Erbb4, Corin*, *Tbx20*, *Fgf12, Fgf13*, and *Pde3a*. Additionally, the low MW treatment had upregulation of *Fgf1*, which has been linked with cardiac regeneration^[63]^.

**Figure 7.**
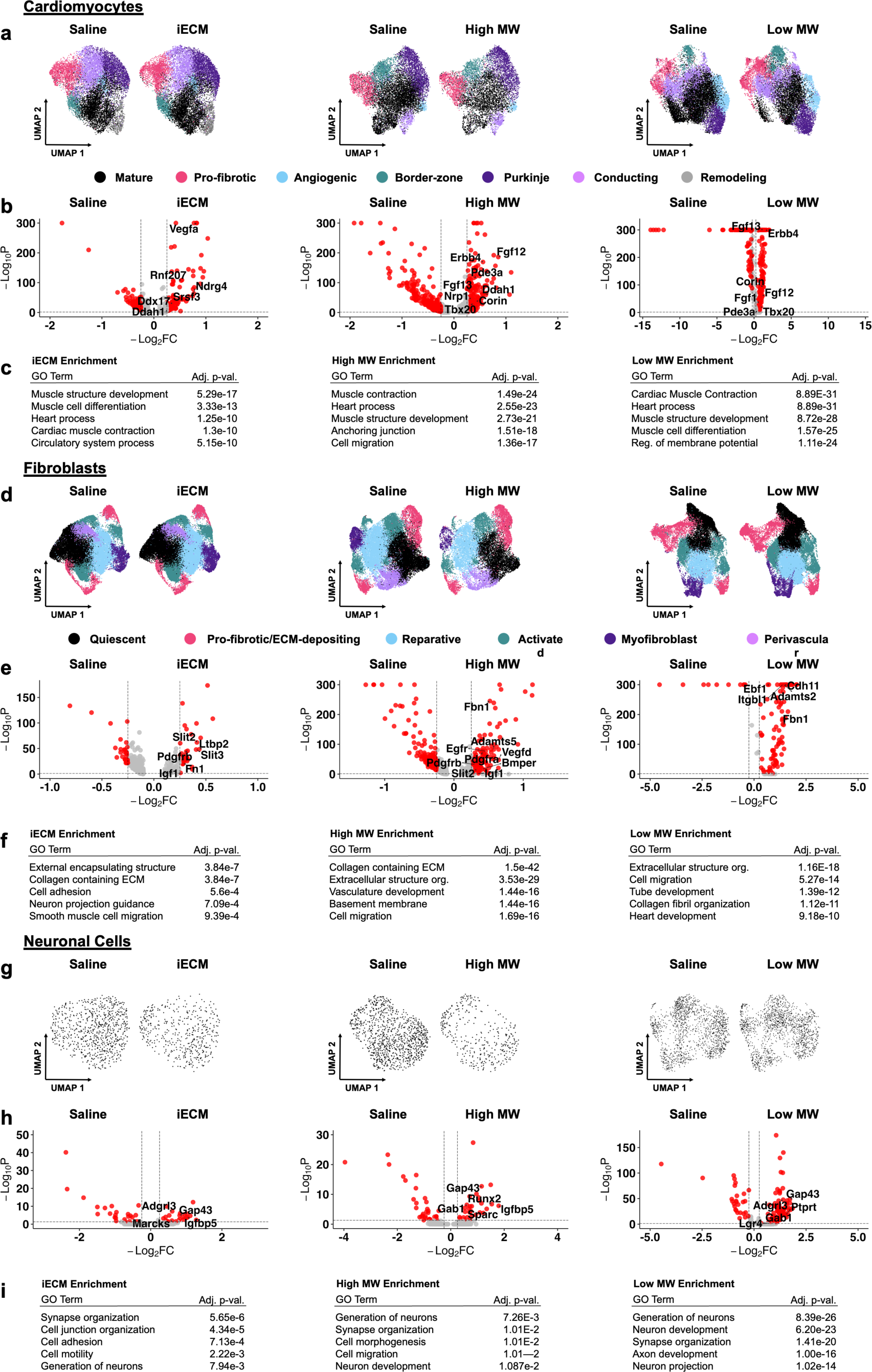
Single nucleus RNA sequencing integration, differential gene expression and gene ontology enrichment analysis of cardiomyocytes for iECM, high MW and low MW compared to saline treatment. The UMAPs, volcano plots and enriched GO terms for iECM, high MW and low MW vs. saline for a-c) cardiomyocytes, d-f) fibroblasts and g-i) neuronal cells. The overall response of the cardiac function related cell types was cardioprotection.

Within the fibroblast subset (*Postn^+^/Dcn^+^*) (**Fig. 7d-f, S12a-c**), DEGs across treatments consisted of genes implicated within fibroblast activation and cardiac fibrosis as well as with cardioprotection (Fig 7a-c). Within the iECM treatment group (Fig 7a), *Slit3*, *Ltbp2* and *Fn1* were upregulated, which has been linked with cardiac fibroblast activation and fibrosis^[64]^. There was also upregulation of *Slit2*, *Igf1* and *Pdgfrb*, which has been linked with reduced fibrosis^[65]^, cardiomyocyte proliferation^[66]^ and reduced inflammation^[48b, 65]^, cardiomyocyte proliferation^[66]^, and reduced inflammation^[48b]^. Within the high MW (Fig 7b), there was similar fibroblast activation^[67]^ with upregulation of *Egfr*, *Itgbl1*, *Fbn1, Vegfd, Adamts5*, and *Bmper* while also showing signs of cardioprotection^[48b, 65-66, 68]^ via upregulation of *Pdgfra, Pdgfrb, Igf1,* and *Slit2*. Within the low MW (Fig. 7c), upregulation of *Cdh11, Fn1*, *Itgbl1* and *Adamts2* pointed towards fibroblast activation^[69]^ while *Ebf1* upregulation pointed towards reduced fibrosis^[70]^. Overall, the response within the fibroblast subsets for all treatments showed signs of cardioprotection.

Within the neuronal cell subset (*Cadm2^+^/Ncam2^+^*) (Figure 7g-i), all treatments showed signs of neuron development and proliferation via *Gap43* upregulation^[71]^. With iECM (**Fig 7g**), upregulated genes included *Marcks,* which is linked with nerve regeneration^[72]^, *Igfbp5* upregulation which is tied to cardiomyocyte survival^[73]^, and *Adgrl3* which has been linked with tissue development^[74]^. With high MW (Fig. 7h), *Igfbp1* upregulation was also seen alongside *Sparc* and *Gab1* which has been linked with fibrosis and cardiomyocyte apoptosis^[75]^. Within the low MW (Fig. 7i), *Gab1* and *Adgrl3* upregulation were seen alongside *Ptprt*^[76]^ and *Lgr4*^[71b]^, which has been linked to neuronal development. Overall, the expected neuronal sprouting genes typical post-MI is seen via *Gap43* expression. In addition, evidence suggests neuronal cell development within all treatments.

#### 2.4.4 iECM and its components exhibit similar gene ontology enrichment

iECM, high MW and low MW exhibited a variety of gene expression that all pointed toward a similar phenotype across all major cell types. When these DEGs were analyzed for gene ontology (GO) pathway enrichment, many conserved pathways were found across the 3 treatments with subtle differences (**Figure 8)**. All treatments enriched for angiogenesis in both endothelial and lymphatic endothelial cells. Within mural cells, all treatments enriched for cell migration suggesting that all the treatments resulted in strong vasculature development signatures. In particular, the high MW had robust evidence for endothelial progenitor cell recruitment within the endothelial cell population, enriched for mesenchymal cell migration within the lymphatic EC population, and enriched for cell junction organization within the mural cells. Within immune populations, all treatments enriched for lymphocyte activation within B and T cells as well as macrophage activation and endocytosis. The fibroblast response was equally similar showing signs of activation through cell adhesion and ECM related pathways. Finally, the pathway enrichment for cardiomyocytes and neuronal cells were consistently pro-repair for all treatments suggesting cardioprotection and the development of neurons.

**Figure 8.**
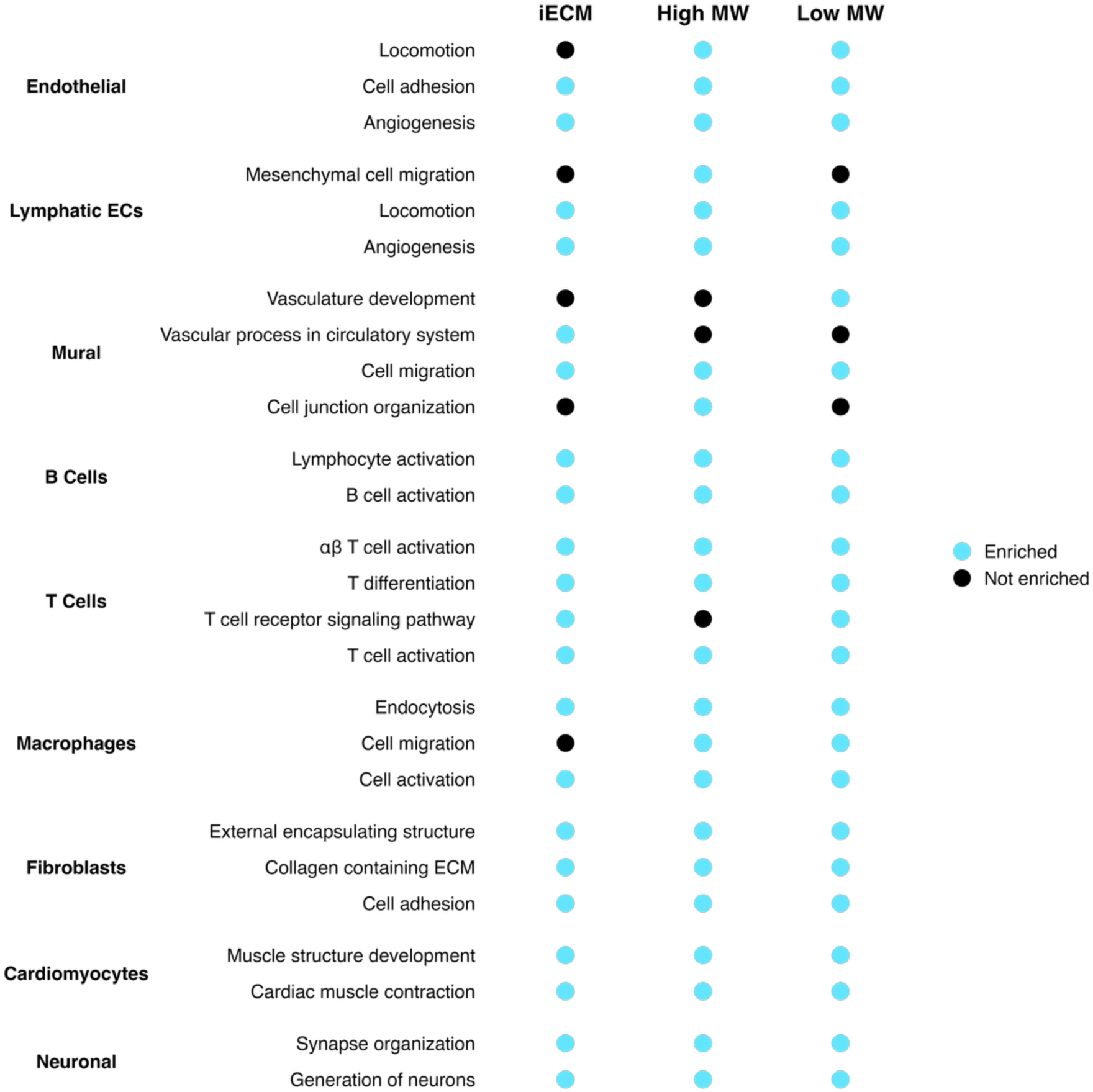
Summary of GO enrichment for each treatment vs. saline per cell type. The enriched pathways were similar between the treatments suggesting an angiogenic response for vascular related cell types, immune cell activation, and pro-reparative phenotypes for cell types related to cardiac function.

The GO summary shows that both components of iECM exhibit pro-reparative effects that are conserved when delivered individually. Additionally, the DEGs were not confined to a single component, indicating that the bioactivity of the high- and low-MW components overlap. When contextualized with the compositional overlap seen in the mass spectrometry results (Fig 2c-d), these data suggest that the differences in morphology and biochemical nature may drive the differences in binding efficiency *in vitro* and *in vivo* while the composition drives the downstream gene expression. However, since the two components do have varying abundances of a similar composition, there still exists subtle differences in the differential gene expression and pathway enrichment analysis, particularly illustrated in the vascular cell type response generated by the high MW. Overall, these gene expression data show that the composition of ECM biomaterials drive the reparative response regardless of their morphology and biochemical features highlighting the intrinsic pro-reparative nature of ECM biomaterials.

## 3. Conclusion

We were able to extensively characterize the material properties and biological mechanisms of action for iECM and its components through methods not previously utilized for ECM biomaterials. Within iECM, we identified high and low MW components that are biochemically and morphologically distinct with differences in *in vitro* binding to ECM components and cellular receptors. Proteomic analysis revealed both components were comprised of peptides from similar proteins, though the relative abundance differed between the two. Evidence from QCM measurements also demonstrated that the high and low MW components interact with one another, further suggesting binding interactions with exposed ECM. These differences seen *in vitro* translate to differences in behavior after administration following MI. While the low MW components alone were able to significantly reduce vascular permeability at the 1-hour timepoint, the high MW components bound to the compromised endothelium more efficiently. However, the combination of both components resulted in increased total signal, suggesting the importance of the total composition of iECM on acute localization. snRNAseq showed that across all major cell types, the angiogenic, immunomodulatory, and pro-regenerative phenotypes were similar across iECM, high MW, and low MW treatment versus saline. Both the DEGs and conservation of GO enrichment between therapeutic treatments groups further highlighted the interrelated behavior of the two iECM components. Collectively, these data suggest that the localization of both iECM components allows them to engage vascular related cell populations, thereby shaping the immune response and impacting cardiomyocytes, key determinants of myocardial function.

While these findings complement current sentiment that ECM biomaterials are therapeutic through their heterogeneity, deep analysis of iECM components and their characteristics reveal new approaches to designing infusible biomaterials. Specifically, this work demonstrates that the flexible nanofibrillar shape combined with multivalent, multitargeted binding as seen in iECM confers high specificity for and retention in vasculature at sites of inflammation. These findings highlight material attributes that can be leveraged for designing effective targeted intravascular biomaterials either via synthetic biomimicry or processing modifications to biologically derived materials.

## 5. Experimental Section

### 5.1:Manufacturing of iECM

Porcine derived LV myocardium was decellularized according to previously published protocols^[11]^. The LV myocardium from whole porcine hearts (Collagen Solutions) was isolated, with the endocardium and epicardium removed. Isolated LV myocardium was ground, passed through a ¼ inch stainless steel grate, and decellularized with 1% sodium dodecyl sulfate (SDS, Fisher Scientific) in phosphate buffered saline (PBS) for 4 days, followed by 24 hours of water rinsing. Material was then lyophilized, milled, and passed through a 60-mesh screen.

Milled decellularized ECM was digested at 10 mg/mL in 1 mg/mL pepsin (Millipore Sigma) in 0.1 M HCl for 48 hours. Following digestion, the solution was neutralized with 1 M NaOH, diluted to 6 mg/mL in 1X PBS and centrifuged at 14000 RCF at 4° C for 45 minutes. The supernatant was collected, dialyzed against pure water for 48 hours, and lyophilized. Dry material was resuspended at 16 mg/mL in PBS, filtered through a 0.22 μm filter, aliquoted, lyophilized and stored at -80° C until use. Prior to use, iECM was resuspended at 10 mg/mL in sterile water and stored on ice until use. iECM components were separated out through resuspension in 50% ethanol, causing the high MW to precipitate out. Solutions were centrifuged at 14000 RCF for 10 minutes to pellet the high MW components. This precipitation and centrifugation method was repeated once. Pellets and supernatants were lyophilized and resuspended with PBS and sterile water, respectively, before use.

### 5.2 :Characterization of iECM Molecular Weight and Relative Composition Analysis

For quantitative molecular weight analysis, iECM was analyzed with size exclusion chromatography (SEC) using a Bio-Rad NGC Quest 10 Plus fast protein chromatography system. 200 μL of 10 mg/mL iECM was flowed at 1 ml/min through at Bio-Rad Enrich SEC 650 column (10 x 300 mm) with a separation range of 5 kDa to 650 kDa. Mobile phase molecules were detected by absorbance at 215 nm. Relative composition of iECM was determined by integrating the area under the curve and determining the relative area of each of the peaks. Apparent molecular weight was determined by constructing a molecular weight curve using a protein gel filtration standard (Bio-Rad) with a range of 670 kDa to 1.35 kDa. Confirmation of iECM fractionation was done using SEC and through SDS polyacrylamide gel electrophoresis (PAGE). A 4-12% Bis-Tris gel (Invitrogen) and a broad range prestained protein ladder (Abcam) were used for PAGE. Protein bands were visualized using an Imperial protein stain (ThermoFisher).

### 5.3 :Characterization of iECM Components

#### 5.3.1 : Biochemical Characterization

Free amines in iECM components were determined using an o-Phthaldialdehyde reagent assay (ThermoFisher) and compared against glycine standards. Free thiols were determined using a thiol detection assay kit (Cayman Chemical) and compared against cysteine standards. Sulfated glycosaminoglycans were determined using a 1,9-dimethylmethylene blue assay and compared against chondroitin sulfate standards^[77]^. dsDNA was isolated using a NucleoSpin kit (Macherey-Nagel) and quantified using a PicoGreen Fluorescent reporter (Invitrogen). To determine fluorophore conjugation efficiency to iECM components, 10 mg/mL complete iECM was labeled with 10 mg/mL Vivotag-S 750 (VT-750, Perkin Elmer) at a volume ratio of 1:100. Labeling was done at room temperature for 1 hour, after which iECM was separated into components as described above and lyophilized. Dried labeled iECM and iECM components were resuspended at their original volumes and their fluorescence was determined with a UV-Vis spectrophotometer plate reader. Fluorophore labeling efficiency was determined by comparing fluorescence of components to the fluorescence of complete iECM.

#### 5.3.2 : Mass Spectrometry Proteomics

iECM was subjected to trypsin digestion using S-Trap micro spin columns (Protifi #C02-micro) according to the manufacturer’s protocol. In brief, 2x S-Trap lysis buffer was added to each sample to reach 1X concentration. Samples were reduced using Tris (2-carboxyethly) phosphine (TCEP), alkylated with 2-chloroacetimide (ClAA), and digested with trypsin (1:100 enzyme:substrate ratio) for approximately 15 hours at 37°C. Digested peptides were desalted using Pierce C18 spin columns (Thermo Fisher #84850) according to the manufacturer’s protocol.

Digested peptides (200 ng) were loaded onto individual Evotips and separated on an Evosep One chromatography system (Evosep, Odense, Denmark) using a Pepsep column, (150 µm inter diameter, 15 cm) packed with ReproSil C18 1.9 µm, 120Å resin. With a “30 samples per day” LC gradient, the sample was analyzed on a timsTOF Pro mass spectrometer (Bruker Daltonics, Bremen, Germany) via the nano-electrospray ion source (Captive Spray, Bruker Daltonics) in PASEF mode. Ramp time was 100 ms and 10 PASEF MS/MS scans per topN acquisition cycle were acquired. MS and MS/MS spectra were recorded from m/z 100 to 1700 and ion mobility scanned from 0.7 to 1.50 Vs/cm^2^. Precursors for data-dependent acquisition were isolated within ± 1 Th and fragmented with an ion mobility-dependent collision energy, which was linearly increased from 20 to 59 eV in positive mode. Low-abundance precursor ions with an intensity above 500 counts and below 20000 counts were repeatedly scheduled and otherwise dynamically excluded for 0.4 min.

Data was searched via MSFragger v4.0 via FragPipe v21.1 against the UniProt protein database restricted to *Sus scrofa* with added common contaminant sequences^[78]^ (46,285 total sequences, downloaded 1/8/25)^[79]^. Precursor tolerance was ±15 ppm and fragment tolerance was ±25 ppm. Enzyme cleavage was semi-specific trypsin. Fixed modifications were carbamidomethyl (C). Variable modifications were oxidation (M), oxidation (P) (hydroxyproline), deamidation (NQ), Gln->pyro-Glu (N-term Q), and acetyl (protein N-terminus). IonQuant v1.10.12 was used for label free quantification with match-between-runs enabled and default parameters. Results were filtered to 1% FDR at the peptide and protein level.

#### 5.3.3 : Transmission Electron Microscopy

First, glow discharged, 200 mesh, Formvar carbon copper TEM grids were inverted onto droplets of iECM, high MW and low MWs for 2 minutes. Grids were washed 3 times for 30 seconds using 0.22 µm filtered double-distilled water and stained with 2% uranyl acetate for 1 minute and finished with a 5-second blot. Samples were air-dried before imaging on a Tecnai G2 Spirit BioTwin (FEI) operated at 80KeV and equipped with a bottom-mounted 4kx4k Eagle camera. Images were collected at 2.9kx, 6.8kx, 11kx, and 30kx.

#### 5.3.4 : Quartz Crystal Microbalance

Frequency and dissipation data was collected under flow conditions on a QCM-D QSense Explorer (Biolin Scientific). Gold coated chips were used (Biolin Scientific QSX 301). Prior to experiments, chips were soaked in 2% sodium dodecyl sulfate in water at 70°C for 2 hours. Chips were washed 5 times with room temperature deionized water, dried and stored in the original boxes. Prior to experiments, chips were cleaned within a UV/ozone cleaning chamber for 10 minutes.

N-hydroxysuccinimide functionalized lipid suspensions (NHS-DPPC) were produced similar to previous reports^[28]^. Equal mass (15 mg) of palmitic acid N-hydroxysuccinimide ester (Sigma P1162) and 1,2-dipalmitoyl-sn-glycero-3-phosphocholine (Sigma P4329) were mixed with 15 mL of 0.1 M Mes [2-(2-morpholino) ethanesulfonicacid] buffer (pH 6.0). Lipid solutions were heated to 60°C and mixed. Lipid suspensions were aliquoted and frozen in -80°C until use.

For binding interaction between high MW and low MW components, the high MW and low MW were coated at a concentration of 5 mg of ECM/mL to ensure consistent exposure of the components during conjugation.

Basal lamina ECM components and endothelial cell surface receptor protein solutions were diluted to 0.1 mg/mL from the stock using 1x phosphate buffered saline (1x PBS). Solutions were aliquoted and frozen at -80°C until use. ECM components used were type I collagen (Advanced BioMatrix 5005100ml 3 mg/mL), type IV collagen (Cultrex Mouse Collagen IV 0.5 mg/vial 3410-010-01), laminin 111 (Gibco 1.2 mg/mL 23017015), and laminin 521 (LA5-H5215-500ug). Cell surface receptor proteins used for conjugation were P-selectin (Acro 50-165-8924 – 100 ug), integrin α1β1 (Acro 50-253-4563 – 100 ug) and ICAM/CD54 (MedChemExpress HY-P7262).

Antibodies used for validation of ECM and cell surface receptor protein conjugation were diluted to 50 ug/mL using 1x PBS. ECM antibodies were type I collagen (Invitrogen PA5-29569), type IV collagen (Invitrogen PA1-85320), laminin 111 (Biosensis R-1808-100) and laminin α5 antibody (Abnova H00003911-D01P). Cell receptor antibodies were P-selectin (Invitrogen 12-0626-82), integrin α1 (Invitrogen MA5-55280), and ICAM/CD54 (Invitrogen 500-P287-100UG).

Flow experiments were carried out at 100 uL/min unless otherwise noted. A 22°C baseline was collected with DI water. NHS-DPPC was introduced for 5 minutes followed by a 5-minute 1x PBS wash. ECM or cell surface receptor proteins were introduced for 10-20 minutes at 50 uL/min for conjugation followed by a 1x PBS wash at 100 uL/min.

For antibody assay validation, the temperature was maintained at 22°C. After the 1x PBS wash following conjugation, a 1 mg/mL bovine serum albumin (BSA) (Geminin Bio 50-753-3052) solution in 1x PBS was used for blocking followed by a 5-minute 1x PBS wash. The antibody was introduced at 25 uL/mL followed by a final 1x PBS wash.

For the binding assays of high MW to low MW and vice versa, following the 1x PBS wash after conjugation, the temperature was changed to 37°C. After equilibrium, BSA was used for blocking followed by a 5-minute 1x PBS wash at 100 uL/min before the high MW or low MW (∼1.9 mg of ECM/mL and ∼8.1 mg of ECM/mL respectively) was flowed at 100 uL/min for 10 minutes followed by a final 1x PBS wash.

For iECM, high MW and low MW components binding experiments, following the 1x PBS wash after target conjugation, the temperature was changed to 37°C. At 37°C equilibrium, BSA was used to block followed by a 5-minute 1x PBS wash at 100 uL/min. Then, the iECM, high MW components or low MW components (10 mg of ECM/mL, ∼1.9 mg of ECM/mL and ∼8.1 mg of ECM/mL) or non-specific peptide controls (∼8.1 mg/mL) of neutral charge poly-L-proline <10 kDa (Sigma P2254-100MG) and positive charge poly-L-lysine <15 kDa (Sigma P6516-500MG) were flowed at 100 uL/min for 10 minutes followed by a final 1x PBS wash.

Frequency and dissipation data was exported from the QCM-D control software as a text file. The data was imported into the NBS-QCMAnalysis software (Nanobiosensorics Laboratory). Baselines were created using equilibrium timepoints, e.g. the 1x PBS wash after BSA blocking. After baselining, the timepoints of interest were isolated, exported and the differences in the equilibrium frequencies for the 3^rd^ overtone were extracted and calculated for semiquantitative comparison. All conditions were performed in triplicate.

### 5.4 Testing iECM and iECM Components in Rat Model of MI and I/R Injury

All procedures in this study were approved by the Committee on Animal Research at the University of California San Diego and in accordance with the guidelines of the Association for the Assessment and Accreditation of Laboratory Animal Care (A3033-01).

#### 5.4.1 : Rat model of MI and I/R Injury

To induce MI, adult female Sprague-Dawley rats (225-250g) were used as previously described^[11, 32]^. To induce acute MI, the left coronary artery was accessed via a thoracotomy and ligated for 35 minutes with a suture. Following ischemia, the suture was released to restore blood flow. For simulated intracoronary infusion, the aorta was clamped and the 200 μL of treatment (saline, 10 mg/mL iECM, 1.9 mg/mL high MW components, 8.1 mg/mL low MW components) was injected into the LV lumen using a 30 G needle. Infusions were performed within 10 minutes of reperfusion.

#### 5.4.2 : Biodistribution of iECM and iECM Components

To determine biodistribution and LV localization of iECM and iECM components, iECM was labeled with VT-750 at a ratio of 1:100 by volume. iECM was labeled prior to fractionation with ethanol to preserve the relative labeling ratio found in complete iECM. Labeled material was delivered in the rat model of MI as previously described^[11]^. At time points of 1 hour and 3 days, rats were euthanized and perfused with PBS to remove residual iECM in the blood. Heart and satellite organs (lungs, kidneys, spleen, liver) were extracted, with the heart sliced into 6 transverse slices. Heart slices and satellite organs were scanned with an Odyssey CLx (LICORbio) near infrared scanner to determine total signal.

#### 5.4.3 : Effects on Vascular Permeability

To determine the effects of iECM components on vascular permeability, iECM and iECM components were delivered in a rat model of MI as described above. At time points of 30 minutes and 3 days post reperfusion, 1 mL of 0.06 mg/mL bovine serum albumin (BSA) labeled with Alexa Fluor 680 (ThermoFisher) was administered intravenously through the tail vein. 30 minutes after delivery, animals were euthanized and perfused to remove residual labeled BSA. Hearts were extracted and sliced into 6 transverse slices. Heart slices were scanned as described above to determine the amount of BSA that extravasated into myocardium using total signal.

#### 5.4.4 : Single Nucleus RNA Sequencing

Rats underwent the MI I/R injury model as described in 5.4.1. Treatment groups included saline, complete iECM, high MW and low MW. Animals were harvested 7 days post-MI. Heart tissue was processed as previously described^[32]^. In short, hearts were cut into 6 transverse sections. The LV free wall from even sections were isolated, chopped and flash frozen in liquid nitrogen and stored at -80°C before nuclei isolation. Mid infarct sections underwent hematoxylin and eosin staining and brightfield imaging (Olympus VS200). Infarcts were quantified using QuPath by (1) tracing the infarct, (2) the LV free wall with septum and not RV free wall, and (3) the LV lumen. The infarct percentage was then calculated. Hearts below 15% were not used.

Nuclei isolation was performed similar to previous studies^[80]^. Heart tissue from 4 animals per treatment were pooled into 2 biological replicate samples. Approximately 60 mg of pooled tissue was resuspended in nucleus lysis buffer (Sigma NUC101) with RNase OUT inhibitor (Invitrogen 10777019). Samples were minced on wet ice using scissors (FST 14058-11) before homogenization with a dounce grinder (Sigma D8938). An additional 1 mL of lysis buffer was added, and samples were incubated for 10 minutes on ice prior to filtration through 100, 50 and 20-μm cell strainers (CellTrics NC1037263, NC9491906, NC9699018). Samples were pelleted at 1,000g for 5 min at 4°C and resuspended in 2 mL of sucrose gradient buffer. Nuclei were pipetted on 4 mL of sucrose gradient buffer before being pelleted at 1000g for 10 min at 4°. The pellet was washed with nucleus storage buffer, pelleted, washed with 5% BSA in 1x PBS, pelleted and resuspended in 200 μL of 2% BSA and 5 μL of DAPI. Nuclei were diluted with trypan blue, loaded onto a hemocytometer (Bulldog-Bio DHC-N21) and intact nuclei were counted using a brightfield phase-contract (Keyence BZ-X).

Nuclei were then subject to the 10x Chromium Universal 3’ Gene Expression v4 protocol. Paired-end sequencing was performed on an Illumina NovaSeq instrument. Low-level analysis, including demultiplexing, mapping to a reference transcriptome and eliminating redundant unique molecular identifiers (UMIs), was performed with the Cell Ranger 9.0.1 pipeline for the 10X samples.

#### 5.4.5 : Quality control, normalization, integration, coarse clustering, subset analysis and GO enrichment analysis

SN RNAseq data was analyzed using methods adapted from previously^[80]^ in addition to the latest version of Seurat 5.2.0^[81]^. After alignment, the counts matrices were adjusted using SoupX. Cells with a minimum of 200 uniquely expressed genes and maximum of 2500 with less than 5% mitochondrial content were isolated. Mitochondria related genes were then removed. The data was normalized using log normalization. Cells were analyzed unintegrated before integrated analysis using IntegrateLayers(), RPCA reduction and a resolution of 0.1 for the coarse clustering. The coarse clusters were identified for general cell type populations and subset for further analysis.

Subset clusters were isolated in groups of two treatments and re-normalized using SCTransform, clustered and subject to FindAllMarkers(). Clusters containing non-endogenous gene markers were contaminants and thus removed. This subset cleaning process was performed iteratively. With cleaned subsets, DEGs were calculated for treatment vs. saline, with adjusted P value < 0.05, log fold change threshold of 0.25, min.pct = 0.25 to define the DEGs. These DEGs were plotted on volcano plots with positive and negative DEGs set as inputs for GO analysis via GSEA. The most relevant pathways were selected and organized by FDR/adjusted P-value.

### 5.5: Statistical Analysis

Statistical analysis was performed on GraphPad Prism10. For analysis of two groups, a two-tailed unpaired t-test with Welch’s correction was performed. For analysis of multiple groups, a one-way ANOVA with post hoc Tukey’s test was performed. For analysis of multiple groups at multiple conditions, a two-way ANOVA with post hoc Tukey’s test was performed. Significance was accepted at p<0.05.

## Author Contributions

Michael Nguyen: Conceptualization, Methodology, Validation, Formal analysis, Investigation, Data curation, Visualization, Writing – Original draft preparation, Writing – Review & Editing, Project administration.

Alexander Chen: Conceptualization, Methodology, Validation, Formal analysis, Investigation, Data curation, Visualization, Writing – Original draft preparation, Writing – Review & Editing, Project administration.

Van K. Ninh.: Investigation, Formal Analysis, Data Collection, Writing – review & editing.

Maxwell C. McCabe: Investigation, Formal Analysis, Data Collection, Writing – review & editing.

Quincy Lyons: Investigation, Formal Analysis, Data Collection, Writing – review & editing.

Benjamin D. Bridgelal: Investigation, Formal Analysis, Data Collection, Writing – review & editing.

Connor Uhre: Investigation, Formal Analysis, Data Collection, Writing – review & editing.

Julian Cheng: Investigation, Formal Analysis, Data Collection, Writing – review & editing.

Kate E. Reimold: Investigation, Formal Analysis, Data Collection, Writing – review & editing.

Selena Cao: Investigation, Formal Analysis, Data Collection, Writing – review & editing.

Kirk C. Hansen: Resources, Supervision, Funding Acquisition.

Kevin R. King: Resources, Supervision, Funding Acquisition.

Karen L. Christman: Conceptualization, Validation, Resources, Writing – Review & Editing, Supervision, Funding Acquisition.

## Acknowledgements

We thank the UC San Diego Electron Microscopy Core (UCSD-CECM hydrogel-EM Core, RRID: SCR_022039) and its director, Guillaume Castillon, Ph.D., for equipment access and technical assistance.

This work was supported by NIH NHLBI (R01HL165232) and the California Instititute for Regenerative Medicine (TRAN1-15291). AC was funded by a NIBIB training grant (T32EB009380) and AHA Predoc Fellowship (24PRE1180449). MN was funded by an NHLBI training grant (T32HL0074444) and an AHA Postdoctoral Fellowship (24POST1242447).

