## Supplemental Information for "An infusible decellularized extracellular matrix material binds to vasculature in infarcted myocardium and induces pro-reparative gene expression following acute myocardial infarction through inherent avidity and bioactive signaling"

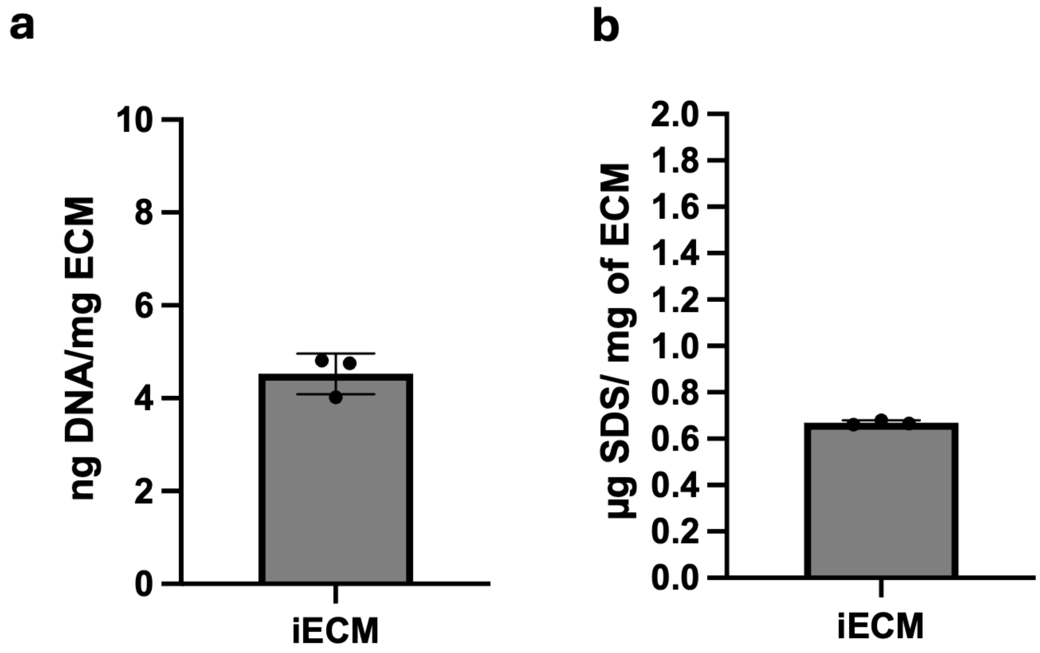


**Figure S1. QC data showing residual double stranded DNA (dsDNA) and residual sodium dodecyl sulfate (SDS).** a) Quantified residual dsDNA within the iECM and its components showing adequate decellularization and b) residual SDS showing sufficient removal of the detergent from after decellularization.

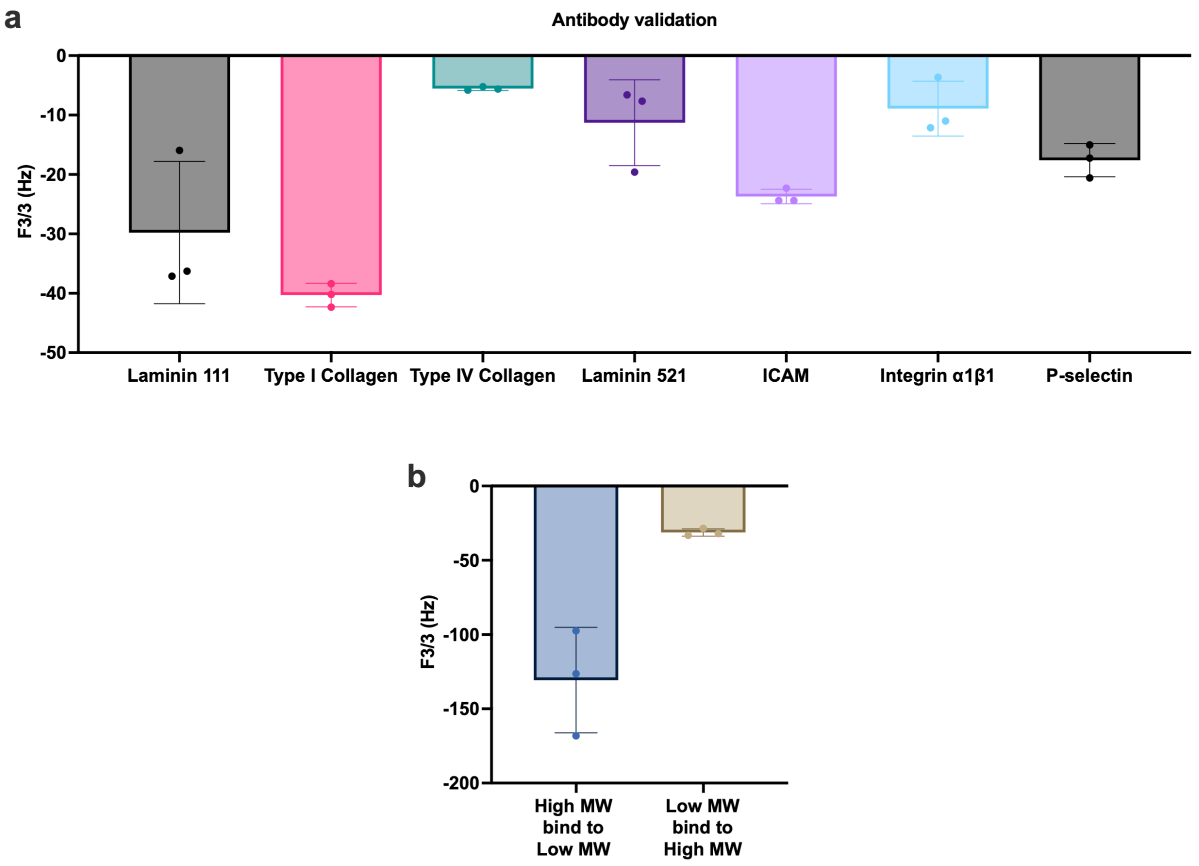


**Figure S2. Assay verification data and interactions between the high MW vs low MW.** The 3rd overtone adjusted frequency shift associated with a) flowing an antibody over the surface of the chemically conjugated ECM proteins and cell surface receptor proteins and b) when one component was conjugated and the other flowed. Third overtone frequency shifts for binding of laminin 111 and type I collagen antibodies were similar to previous reports of antibody binding to ECM coatings via QCM.

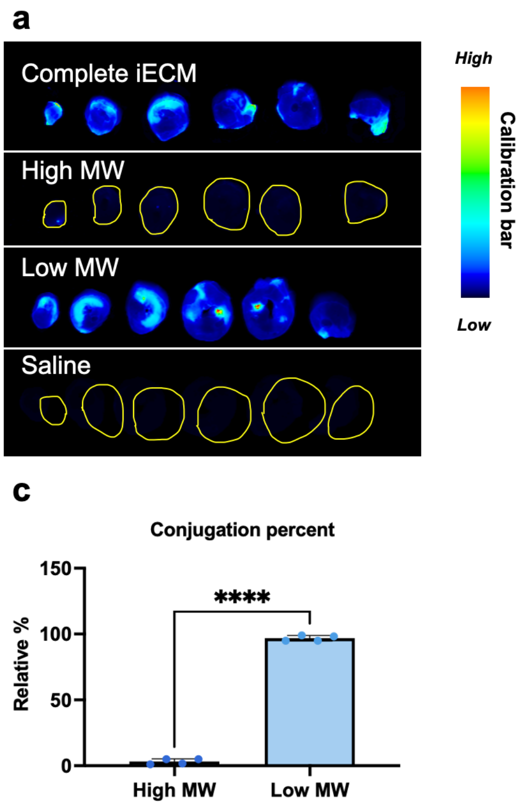


**Figure S3. iECM and components retention and conjugation efficiency.** a) Near infrared imaging of transverse heart sections demonstrating left ventricular localization of complete iECM and iECM fractions. b) Dynamic range adjusted image for high MW components vs saline. c) Conjugation efficiency of the fluorophore for high and low MW components. Welch's two-tailed unpaired t-test **** = $p < 0.0001$.

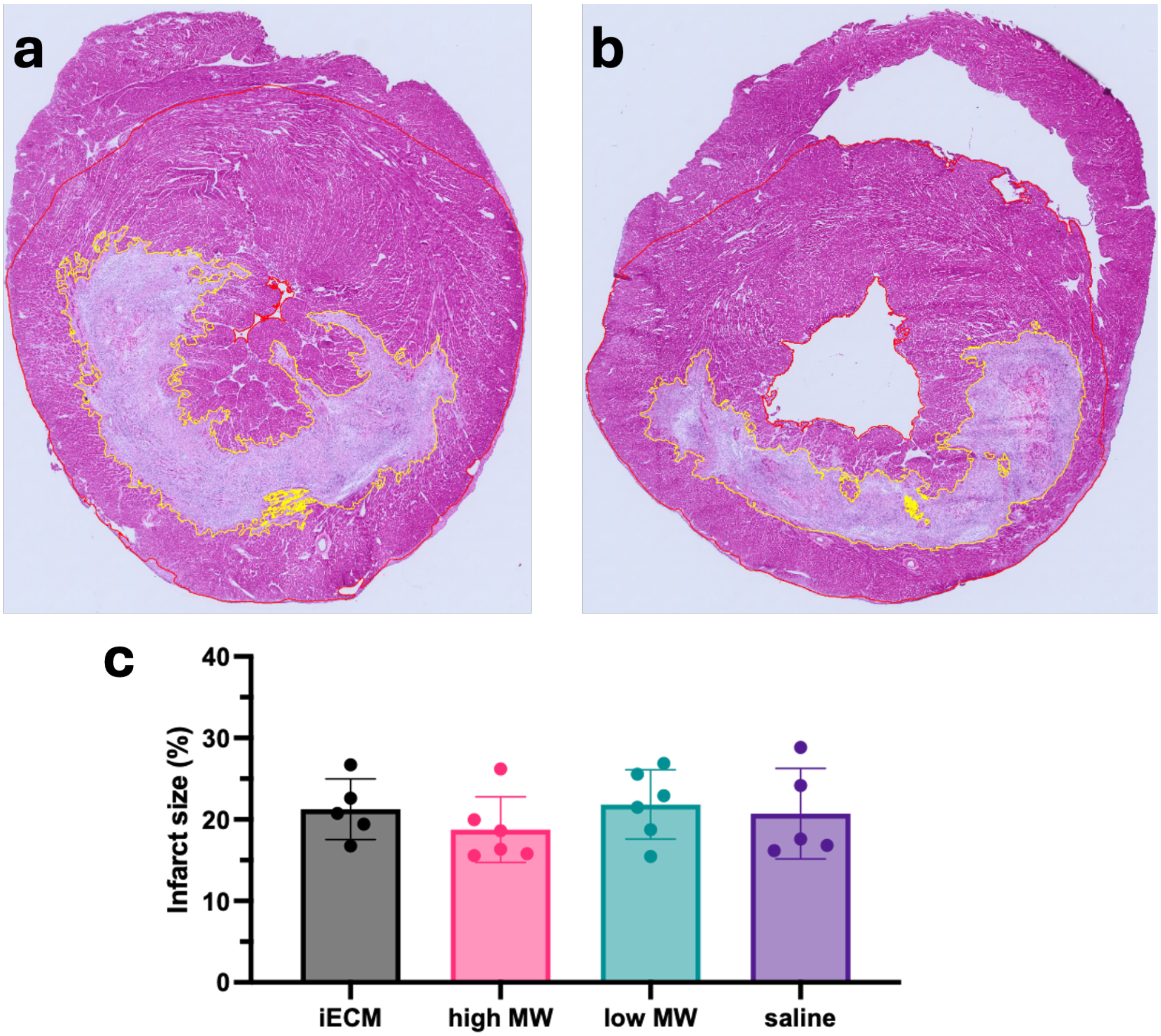


**Figure S4. SN sample QC prior to nuclei isolation and sequencing preparation.** (a-b) Representative H&E images of infarct quantification strategy displaying the three ROIs used for infarct percentage quantification and exclusion. c) Bar graph showing the distribution of quantified infarcts included for each treatment.

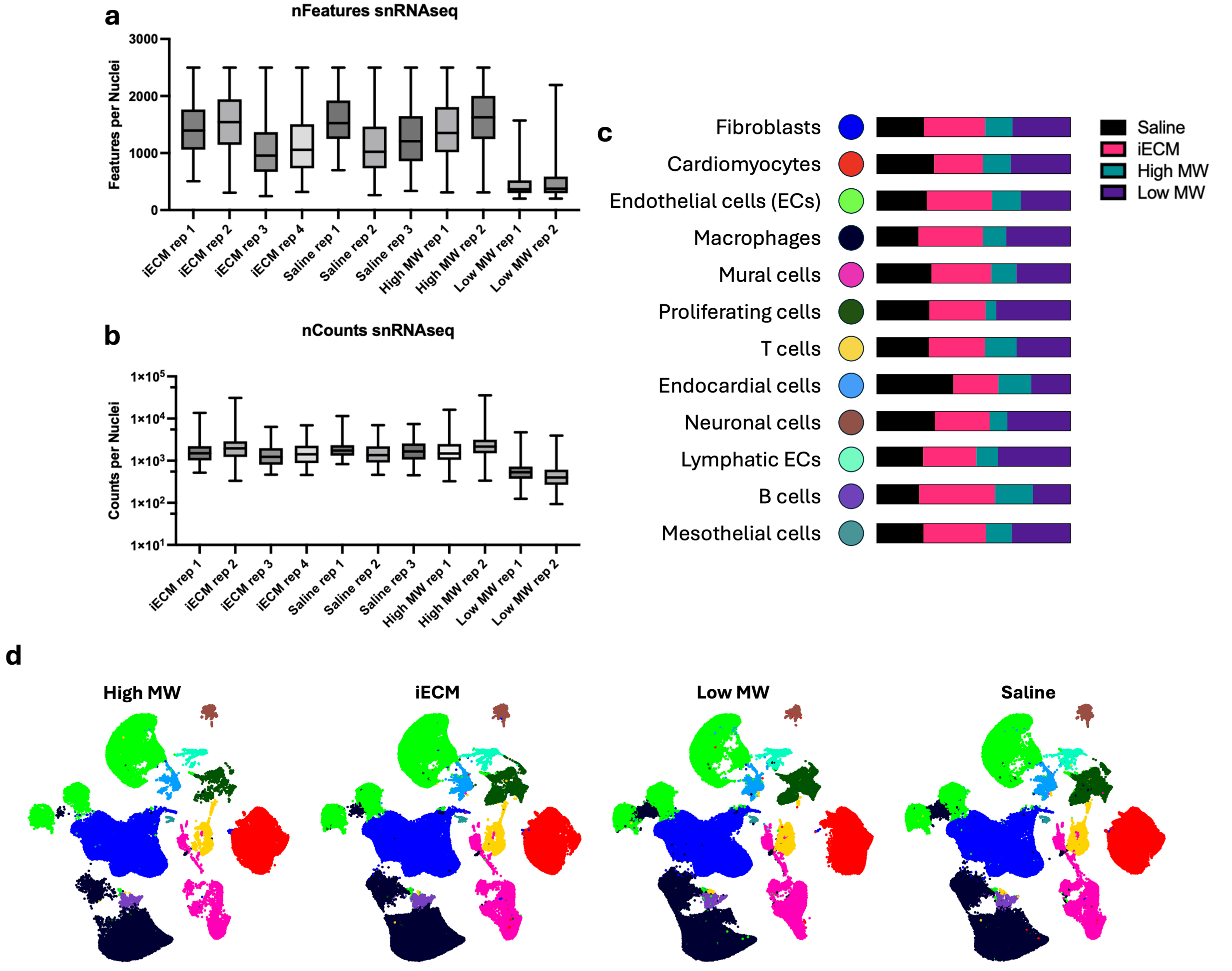


**Figure S5. Single Nucleus RNA Sequencing Quality Control and Cell Type Overview.** a-b) Quality metrics of samples and replicates for single nucleus RNA sequencing samples via features per nuclei (nFeatures) and genes per sample (nCounts). Significant ambient RNA contamination existed within the low MW replicates therefore reducing the features. However, the counts for the low MW replicates were similar suggesting a preservation of the gene expression signal after ambient RNA cleaning. c) The relative percentages of cell for each cell type identified per treatment and the d) associated UMAPs. Sample size n=8 animals pooled into n=4 sequencing replicates for iECM, n=6 animals pooled into n=3 sequencing replicates for saline and n=4 animals pooled into n=2 sequencing replicates for high MW and low MW. Total cells captured: iECM = 151933 cells, low MW = 146616 cells, high MW = 66734 cells, and saline = 125954 cells.

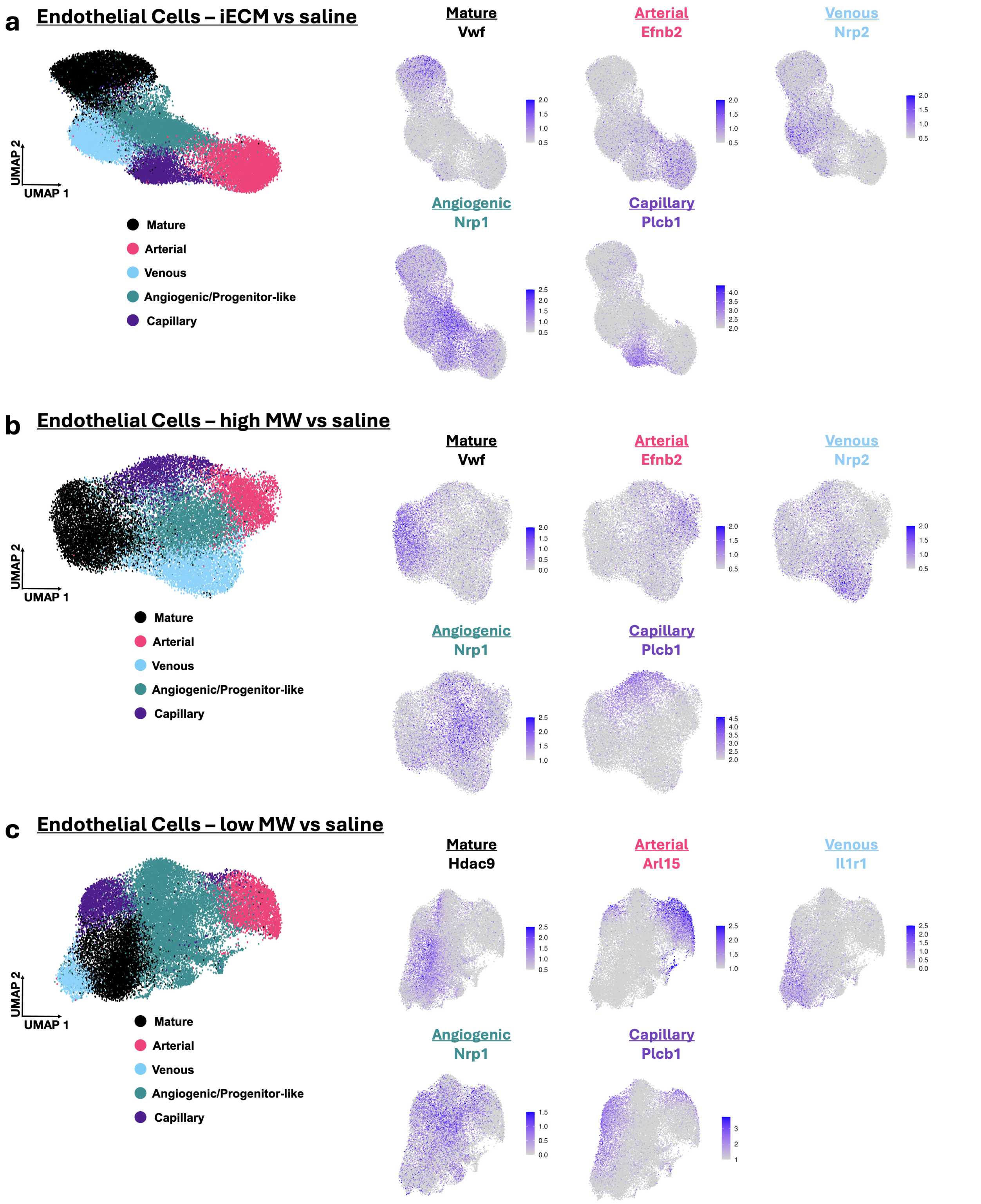


**Figure S6. Breakdown of endothelial cell subpopulations with feature plots of representative genes.** The subpopulations of endothelial cells for a) iECM vs saline, b) high MW vs saline and c) low MW vs saline.

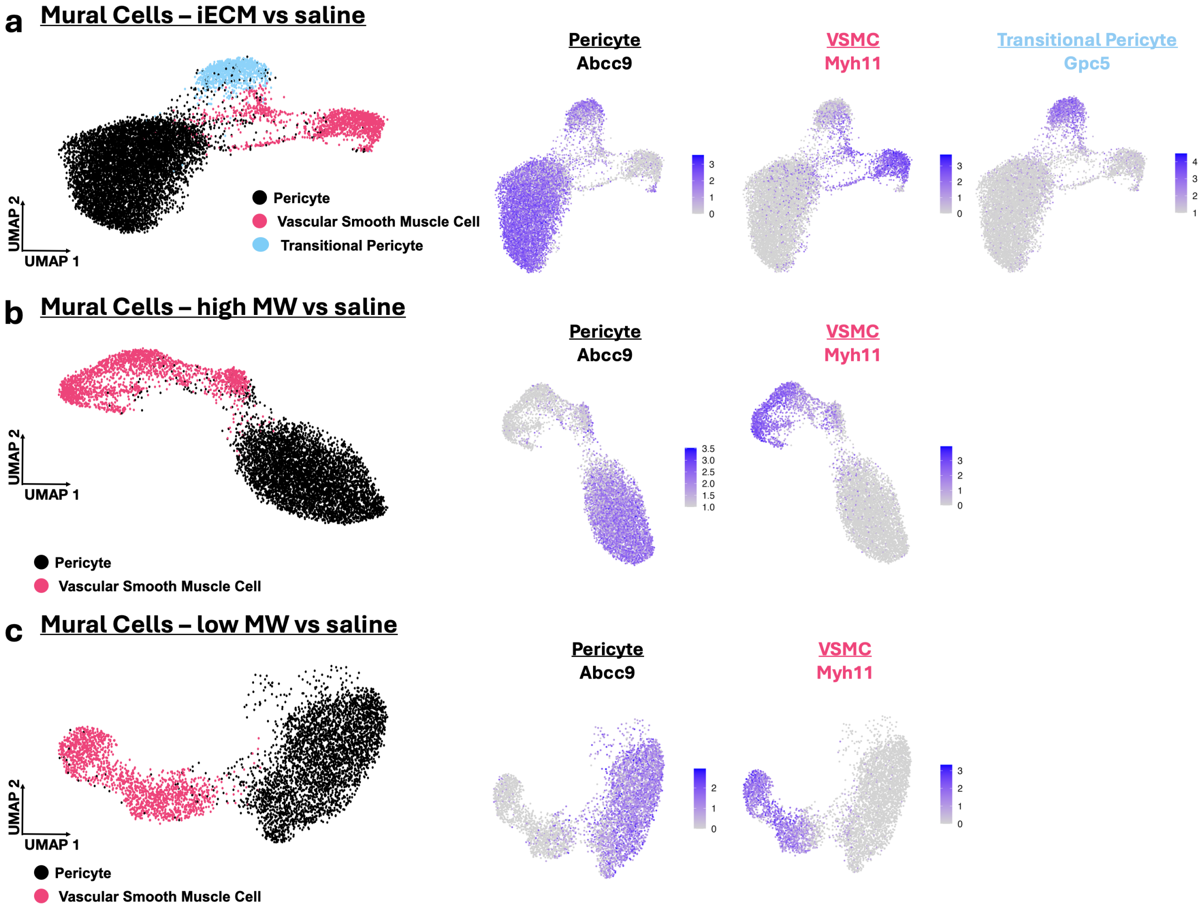


**Figure S7. Breakdown of mural cell subpopulations with feature plots of representative genes.** The subpopulations of endothelial cells for a) iECM vs saline, b) high MW vs saline and c) low MW vs saline.

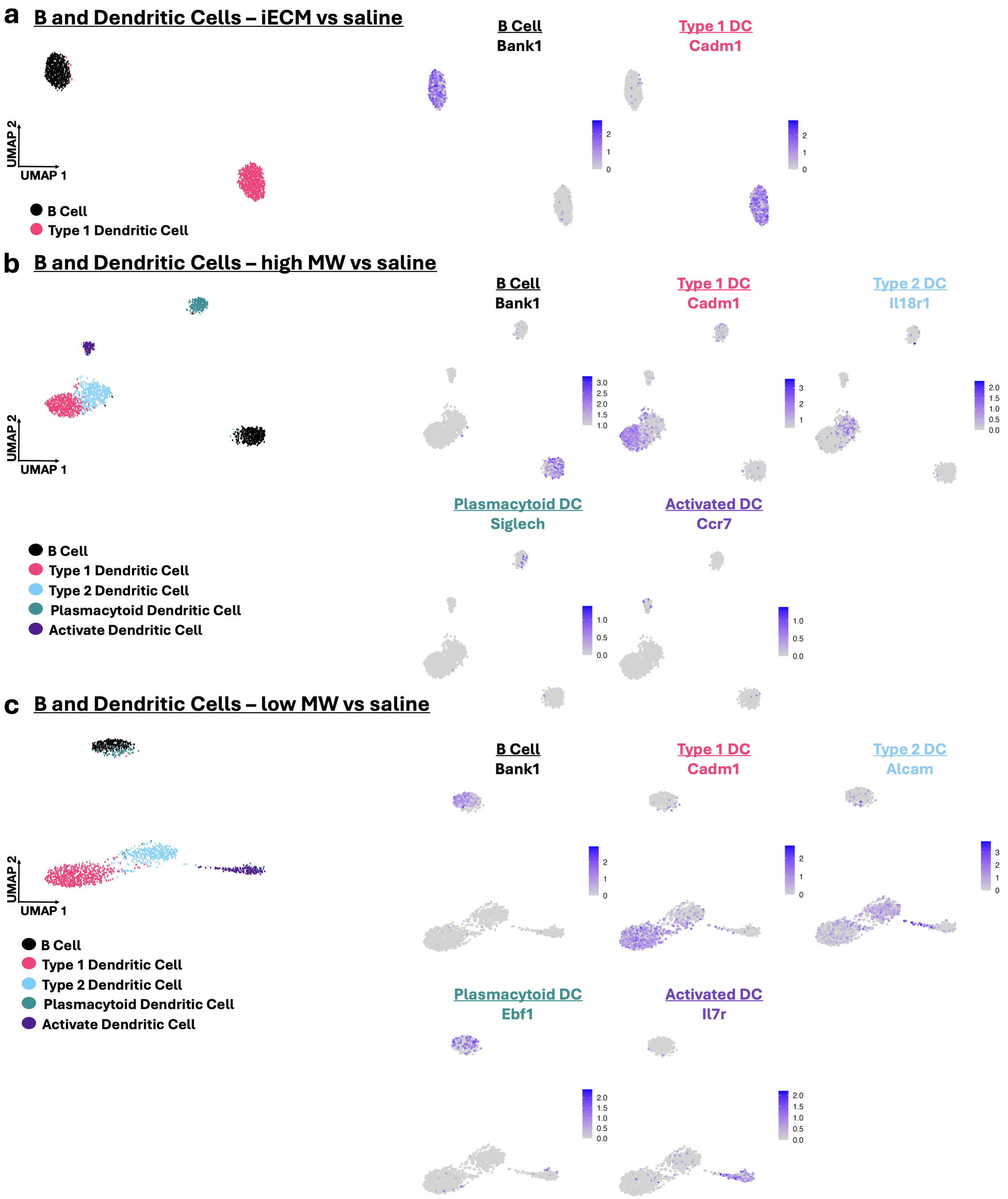


**Figure S8**. **Breakdown of B and dendritic cell subpopulations with feature plots of representative genes.** The subpopulations of endothelial cells for a) iECM vs saline, b) high MW vs saline and c) low MW vs saline.

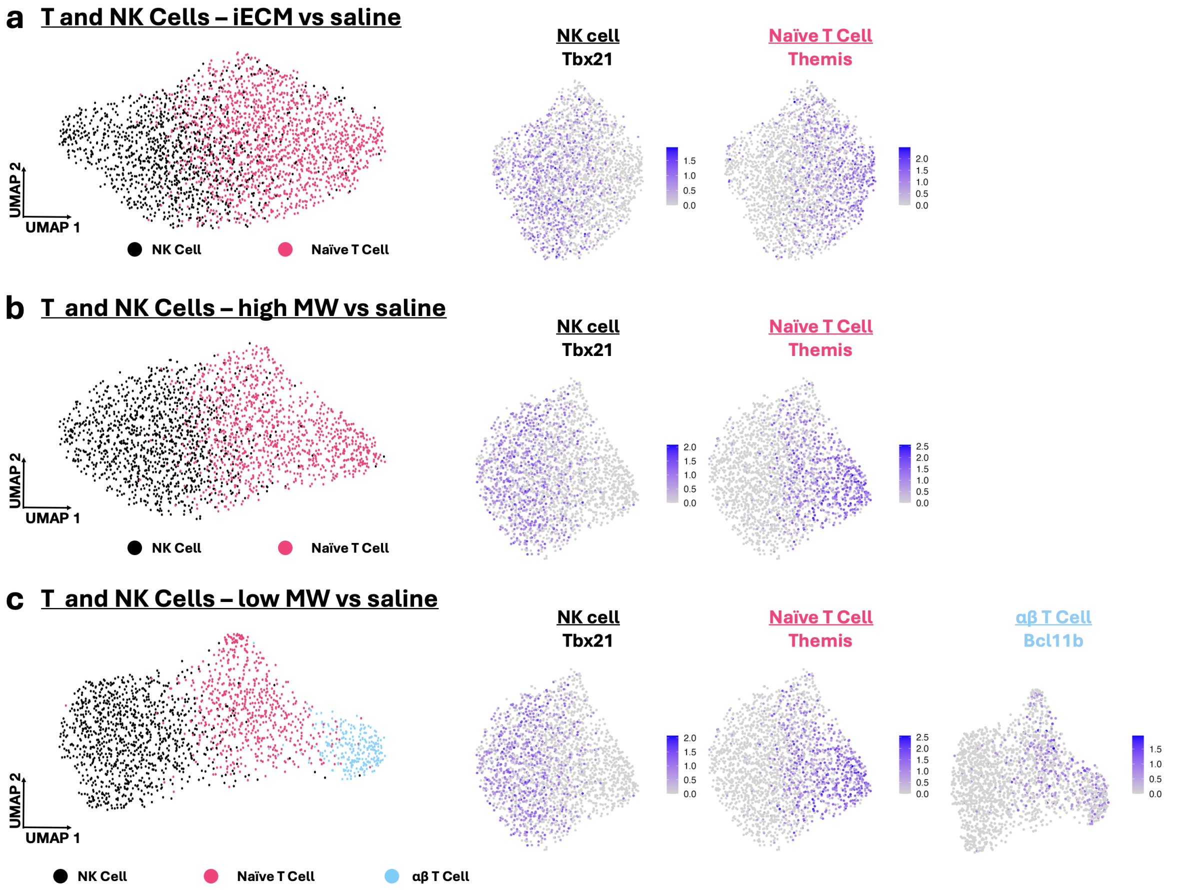


**Figure S9. Breakdown of T and NK cell subpopulations with feature plots of representative genes.** The subpopulations of endothelial cells for a) iECM vs saline, b) high MW vs saline and c) low MW vs saline.

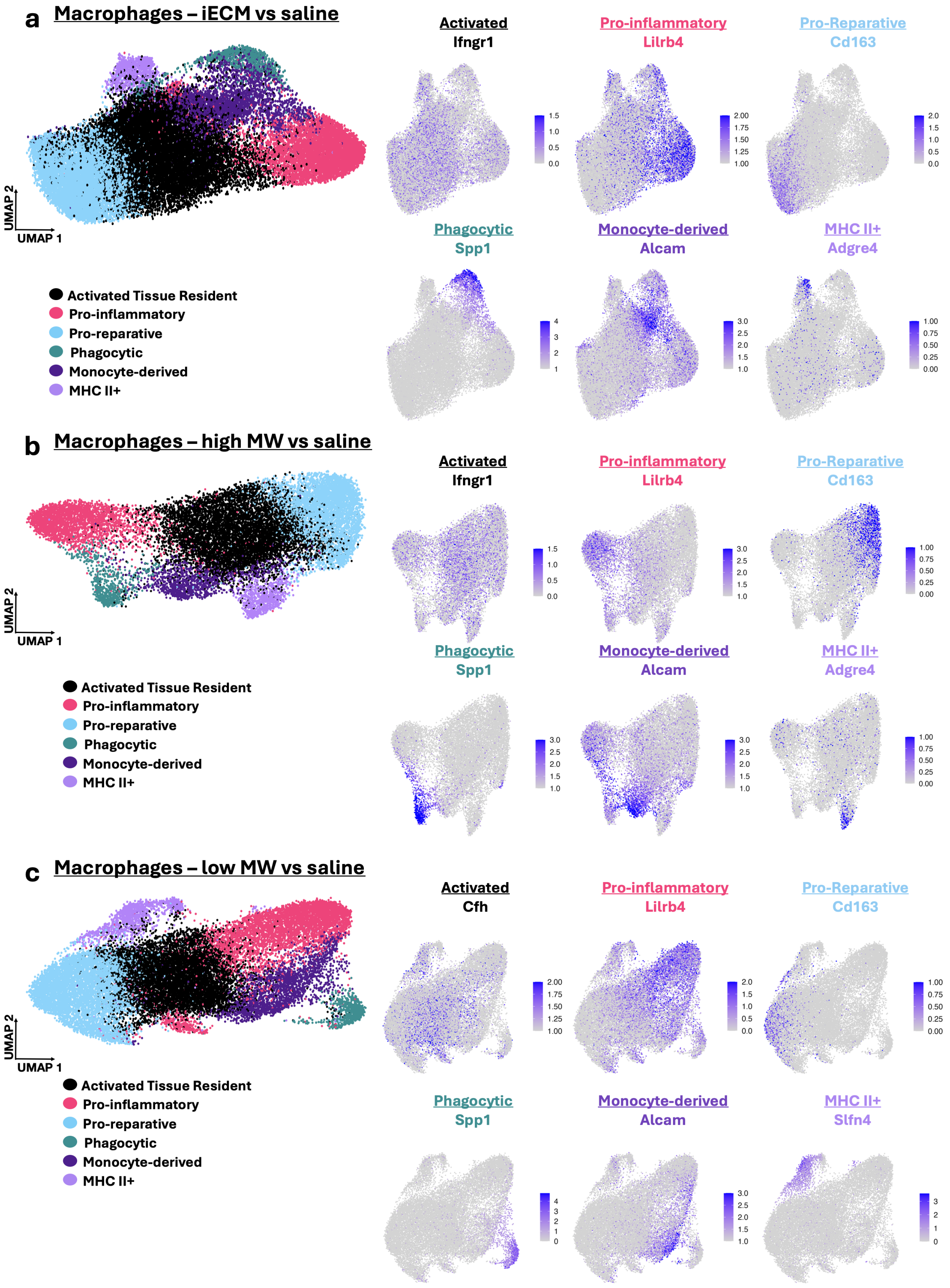


**Figure S10. Breakdown of macrophage subpopulations with feature plots of representative genes.** The subpopulations of endothelial cells for a) iECM vs saline, b) high MW vs saline and c) low MW vs saline.

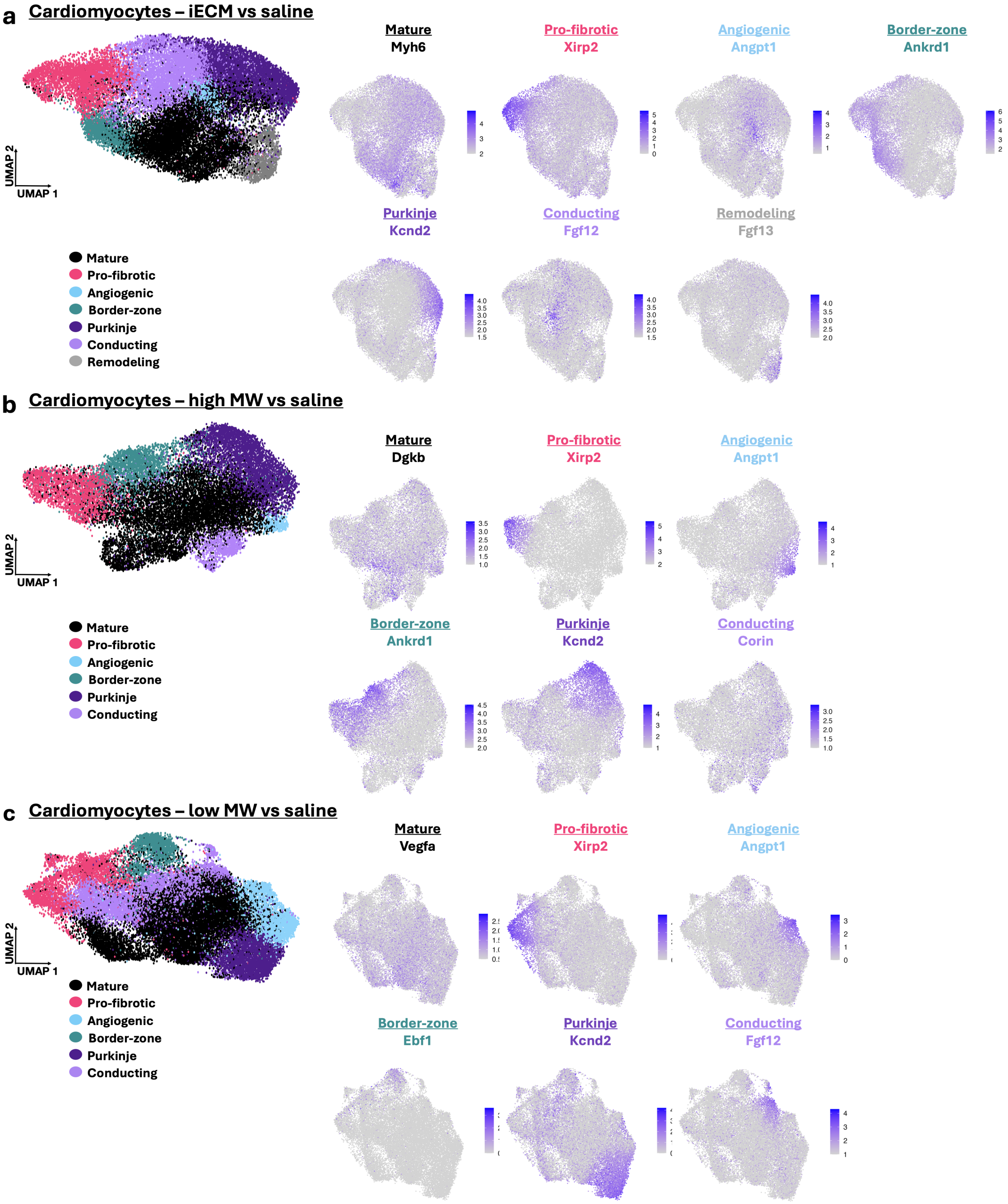


**Figure S11. Breakdown of cardiomyocytes subpopulations with feature plots of representative genes.** The subpopulations of endothelial cells for a) iECM vs saline, b) high MW vs saline and c) low MW vs saline.

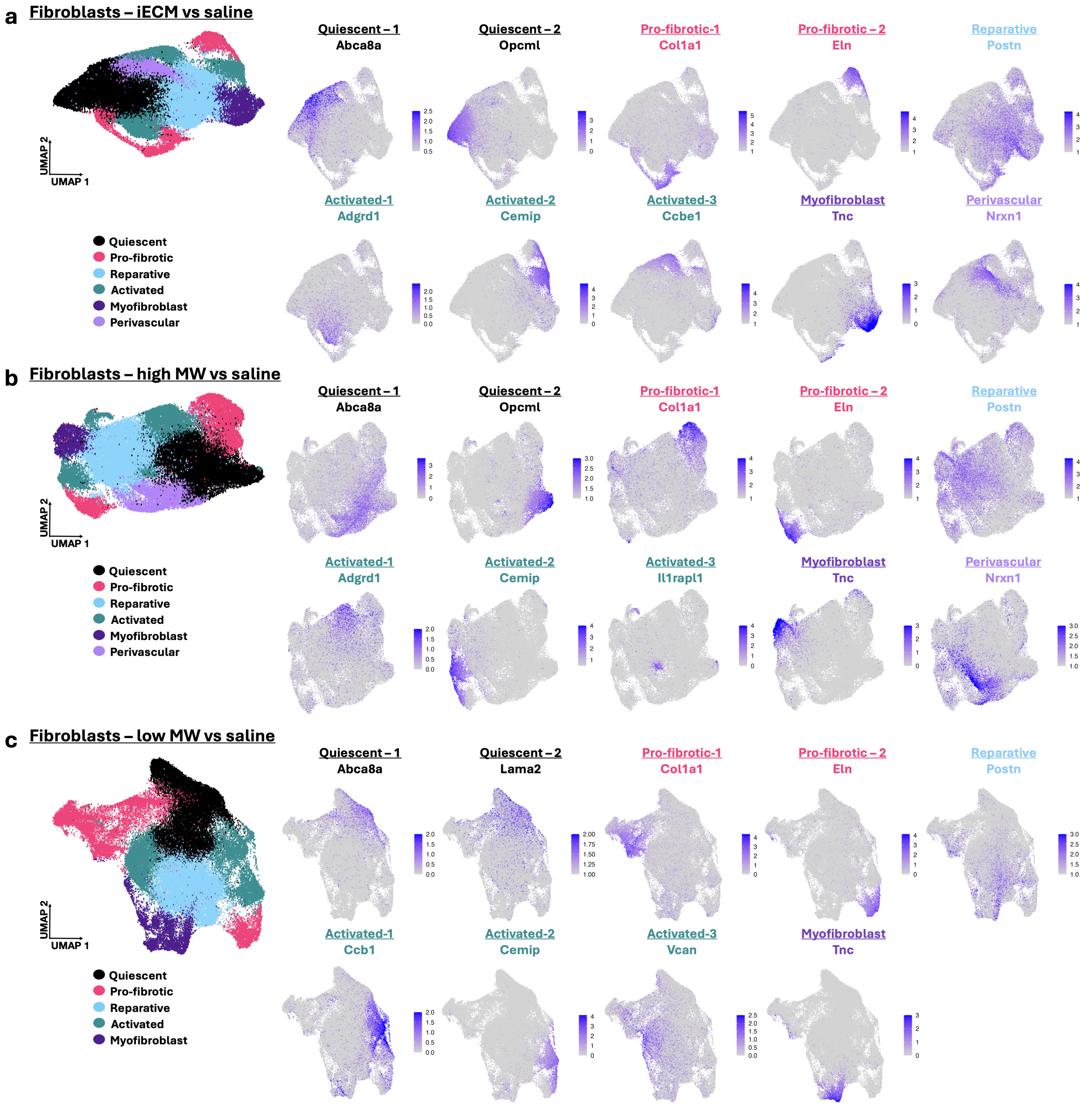


**Figure S12. Breakdown of fibroblast subpopulations with feature plots of representative genes.** The subpopulations of endothelial cells for a) iECM vs saline, b) high MW vs saline and c) low MW vs saline.
